# Uncovering a stability signature of brain dynamics associated with meditation experience using massive time-series feature extraction

**DOI:** 10.1101/2023.06.23.546355

**Authors:** Neil W Bailey, Ben D. Fulcher, Bridget Caldwell, Aron T Hill, Bernadette Fitzgibbon, Hanneke van Dijk, Paul B Fitzgerald

## Abstract

Previous research has examined resting electroencephalographic (EEG) data to explore brain activity related to meditation. However, previous research has mostly examined power in different frequency bands. Here we compared >7000 time-series features of the EEG signal to comprehensively characterize brain activity differences in meditators, using many measures that are novel in meditation research. Eyes-closed resting-state EEG data from 49 meditators and 46 non-meditators was decomposed into the top eight principal components (PCs). We extracted 7381 time-series features from each PC and each participant and used them to train classification algorithms to identify meditators. Highly differentiating individual features from successful classifiers were analysed in detail. Only the third PC (which had a central-parietal maximum) showed above-chance classification accuracy (67%, *p*_FDR_ = 0.007), for which 405 features significantly distinguished meditators (all *p*_FDR_ < 0.05). Top-performing features indicated that meditators exhibited more consistent statistical properties across shorter subsegments of their EEG time-series (higher stationarity) and displayed an altered distributional shape of values about the mean. By contrast, classifiers trained with traditional band-power measures did not distinguish the groups (*p*_FDR_ > 0.05). Our novel analysis approach suggests the key signatures of meditators’ brain activity are higher temporal stability and a distribution of time-series values suggestive of longer, larger, or more frequent non-outlying voltage deviations from the mean within the third PC of their EEG data. The higher temporal stability observed in this EEG component might underpin the higher attentional stability associated with meditation. The novel time-series properties identified here have considerable potential for future exploration in meditation research and the analysis of neural dynamics more broadly.

## 1. Introduction

Mindfulness meditation is a practice that involves focusing attention on the present moment while maintaining non-judgemental awareness of the sensations, thoughts or emotions that arise (Kabat-Zinn, 1994). The practice has been increasingly integrated into clinical practice and is often used across society to improve wellbeing (Baer, 2003; Cramer et al., 2016; La Torre et al., 2022). There is also evidence that mindfulness can improve certain cognitive functions, which suggests a capacity to induce robust changes in brain activity (Bailey et al., 2020; Chiesa et al., 2011; Chiesa & Serretti, 2010; Gill et al., 2020; Im et al., 2021). The identification of alterations to brain activity from meditation poses several potential benefits. If specific patterns of neural activity are altered by meditation practice, then these patterns could be assessed as potential mechanisms by which mindfulness improves mental health (Holzel et al., 2011). Interventions could then be designed to specifically target the patterns of neural activity reflecting mechanisms of action, leading to more effective interventions (Britton et al., 2018; Scangos et al., 2023). The neural activities could also be measured during interventions to determine if the practice is working for a specific individual (Scangos et al., 2023). Individuals who show impairments in the specific neural activities that are enhanced by mindfulness could also be recommended for mindfulness interventions. Finally, the findings related to neural mechanisms of improved mental health from mindfulness could be extended to other fields. For example, targeted neuromodulation using brain stimulation or pharmacological interventions could attempt to replicate the mindfulness-based changes to the neural mechanisms with the aim to improve mental health in individuals who find mindfulness practice prohibitively difficult.

While much research has reported differences in measures of neural activity associated with meditation, findings are inconsistent (Boccia et al., 2015; Falcone & Jerram, 2018; Ganesan et al., 2022; Lomas et al., 2015; Osborn et al., 2022). The inconsistencies relate to both the measures and brain regions in which changes in activity are detected (Lee et al., 2018; Lomas et al., 2015; Schoenberg & Vago, 2019) and in whether neural activity is enhanced or reduced (Lehmann et al., 2012; Osborn et al., 2022). Our own research also follows this pattern, where, depending on the cognitive task or brain regions of neural activity being measured, we have found larger neural responses in experienced meditators (Bailey et al., 2023a), found reduced activity (Bailey et al., 2020), have found altered distributions of activity (Bailey et al., 2020; Bailey et al., 2019a), and also reported no differences compared to non-meditators (Bailey et al., 2019b; Payne et al., 2020). We suspect these inconsistencies might be in part because the effect of meditation is to alter attention processes that underlie the performance of cognitive activities, rather than providing effects that are specific to a particular cognitive domain. As such, different cognitive tasks and experimental conditions may produce different patterns of differences associated with meditation (Wang et al., 2020). However, there may also be deeper mechanistic commonalities underlying the superficial differences across different tasks and experimental conditions. Given the attention training aspect of mindfulness, one potential candidate that has not yet been examined in detail is more stable neural activity, which could underpin increased attentional stability (Bailey et al., 2023b; Lutz et al., 2009). Many other candidate mechanisms are also plausible. To detect potentially novel mechanisms of meditation, this study applied a data-driven approach, using a comprehensive list of over 7000 features extracted from different types of time-series analysis methods to provide the most comprehensive characterisation to date of the time-series patterns in electroencephalography (EEG) data of experienced meditators. Many such time-series methods have not previously been used to study neural activity in meditators. This approach enabled us to determine which features of the data are best for detecting differences in brain activity between experienced meditators and non-meditators, and whether any time-series features of the EEG data may have been overlooked by previous research.

We used resting-state EEG activity instead of a cognitive task, as any cognitive task we selected to study might detect brain activity differences that are only specific to a brain region, network, or function activated to fulfil a specific cognitive process. In contrast, spontaneous resting-state activity reflects ‘baseline’ neural activity that would frequently be engaged in daily life and might therefore be expected to be representative of the neural activity associated with an individual’s daily conscious experience (in contrast to brain activities related to specific cognitive processes, which might only be activated intermittently). Similarly, we did not use meditation-state related data, which can only provide information on state differences that cannot be disentangled from the meditation practice related trait differences of interest, and as such may be less informative of the intrinsic differences in neural activity in meditators (Cahn & Polich, 2006; Lutz et al., 2007).

Research using resting-state EEG to examine the effects of meditation on brain activity has typically examined the power in different frequency bands, which is typically assumed to indicate the strength of neural oscillations. These studies show associations between meditation and altered band power within specific canonical frequency bands: theta (4-8 Hz), alpha (8-13 Hz), beta (13-30 Hz), and gamma (30-80 Hz); with increases in theta and alpha frequency bands most commonly reported (Berkovich-Ohana et al., 2011; Braboszcz et al., 2017; B. R. Cahn et al., 2010; Kerr et al., 2011; Lagopoulos et al., 2009; Lomas et al., 2015; Lutz et al., 2004; Rodriguez-Larios et al., 2021; Wong et al., 2015). Investigations of these frequency bands in healthy non-meditators during the performance of cognitive tasks have indicated that power in each of these frequency bands are associated with a range of functions (Cavanagh & Frank, 2014; Cavanagh & Shackman, 2015; Cooper et al., 2003; Foxes & Synder, 2011; Grunwald et al., 1999; Jensen & Mazaheri, 2010; Kamiński et al., 2012; Kirschfield, 2005; Klimsech et al., 1997; Klimsech et al., 2007; Klimsech et al., 2005). As such, research into frequency band power in meditators has been informative of the effects of meditation on specific brain functions. In keeping with research examining neural activity associated with meditation more broadly, however, findings from studies focused on frequency band power in meditators are also inconsistent (Lee et al., 2018; Lomas et al., 2015; Schoenberg & Vago, 2019).

Furthermore, while neural oscillations are a prominent feature commonly detected in ongoing brain activity, their analysis is only a ‘narrow lens’ through which to characterize the rich variety of temporal patterns that can be observed in general time-varying systems such as the brain (Schoenberg & Vago, 2019). As such, the focus on frequency band power analyses in meditation research has led to an understanding of the effects of meditation on brain activity that is related to changes in neural oscillations, an understanding which may be both over-generalised and incomplete (Schoenberg & Vago, 2019). As such, previous research may have excluded the detection of alternative features of neural activity that may be common across paradigms and may relate more directly to the putative mechanisms of action of mindfulness. For example, quantifying total power within canonical frequency bands commonly involves representing the time-series in terms of a Fourier power spectrum, a representation that captures linear structure in the data. This type of analysis cannot detect nonlinear structure and assumes stationarity (i.e., that statistical properties of the process do not vary over time). As a result, the Fourier power spectrum analysis approach cannot capture non-stationarity (Walker, 1997). However, many other analysis techniques can assess the stationarity of the data (Horváth et al., 2014; Manuca & Savit, 1996; Witt & Kurths, 2002). These analysis techniques may be good candidates to assess neural stability, which we noted earlier may be a mechanism underlying the increased attentional stability reported to be associated with meditation (Bailey et al., 2023b; Lutz et al., 2009).

It is worth noting that there have been a relatively small number of studies of meditation-related brain activity that have explored characteristics of EEG data beyond measures of power in specific frequency bands. These include assessment of the slope of the power-frequency spectrum of aperiodic (non-oscillatory) activity (Bailey et al., 2020) and the nonlinear dynamic complexity and entropy of EEG time series (Aftanas & Golocheikine, 2002; Armbuster-Genc et al., 2016; Vivot et al., 2020; Vyšata et al., 2014). However, all meditation studies we are aware of have still used hypotheses-driven approaches, which examine a small number of manually selected time-series statistics and require subjective and non-systematic choices made by the researcher. The comprehensive analysis methods applied in the current study, which include new (previously untested) types of analysis methods might reveal new insights that are unlikely to be derived purely from conceptual and theoretical perspectives.

To investigate this possibility, we used a comprehensive time-series analysis approach, comparing over 7000 statistical features from the EEG time-series data of experienced meditators and non-meditators. This ‘highly comparative’ approach overcomes the limitations of subjectively selected, small-scale, hypothesis-driven comparisons by systematically searching a comprehensive range of time-series features for the features that best differentiate two labelled groups. Within our study, this ‘highly comparative’ approach was implemented using the highly comparative time-series analysis (*hctsa*) software (Fulcher & Jones, 2017). *hctsa* computes time-series features that assess the linear correlation-based statistics commonly used in EEG research, as well as many other types of features, with a non-exhaustive list including measures of the predictability, stationarity, and self-similarity of the data using entropy, autocorrelation, and fractal scaling developed in fields ranging from seismology to economics, as well as many other features that are not typically assessed within neuroscience (Fulcher & Jones, 2017; Fulcher et al., 2013). This highly comparative approach has previously been used to address a range of questions using EEG data. For example, the approach has been used to extract a data-driven categorization of sleep stages from EEG data and detect the higher order features that separate them (Decat et al., 2022). The approach has also been used to predict individual response to transcranial magnetic stimulation treatment of depression (Bailey et al., 2023e), and to distinguish electrographic seizures from resting brain activity (Fulcher et al., 2013).

Given this background, our aim was to determine whether the EEG data of experienced meditators contains different time-series properties to non-meditators, such that meditators could be accurately identified from their resting-state EEG data. This was achieved by applying a simple classification model using the *hctsa* time-series feature set. If meditators could be accurately identified from the EEG data, our aim was to then characterise the types of time-series properties that best differentiated meditators from non-meditators.

## 2. Methods

### 2.1 Participants

Eyes-closed resting-state EEG data were obtained from two separate studies of experienced meditators, which included a total of 98 participants. After exclusions for low EEG data quality (described in the procedures section), the analysed sample included 95 participants (46 healthy non-meditators and 49 experienced meditators, ranging in age from 19 to 64 years). The results of analyses of the task-based EEG data from these studies have already been published in Bailey et al. (2020); Bailey et al. (2019a); Bailey et al. (2019b); Wang et al. (2020). Participants in these studies were recruited through community advertisements and via meditation centres. Participants in the meditation group were required to have a meditation practice that typically included at least two hours a week of practice, and to have at least six months of meditation experience. Their practice was required to be mindfulness-based with a focus on the breath or body, and for their practice to meet Kabat–Zinn’s definition of “paying attention in a particular way, on purpose, in the present moment, and non-judgmentally” (Kabat-Zinn, 1994). Non-meditators were required to have less than two hours of total lifetime experience with any kind of meditation. All participants were interviewed with the MINI International Neuropsychiatric Interview (MINI) DSM-IV (Sheehan et al., 1998). Potential participants from either group were excluded if they reported any current or previous psychiatric or neurological illness, or current psychoactive medication or recreational drug use. Data from one meditator was excluded due to a history of mental illness. Data from one non-meditator was excluded due to a previous history of meditation practice, and from another non-meditator due to poor data quality (details provided in the procedures section). Trait mindfulness was assessed using the Five Facet Mindfulness Questionnaire (FFMQ) (Baer et al., 2006). The groups did not statistically differ in age, gender, handedness, or years of education (*p* > 0.10, see Table 1), but meditators showed higher FFMQ scores (*p* < 0.001). The study was approved by the ethics committees of the Alfred Hospital and Monash University, and all participants gave written informed consent.

**Table 1.** Self-report data means and standard deviations from each group.

|  | Meditator Group | Non-meditator Group | Statistics |
| --- | --- | --- | --- |
|  | Mean (SD) | Mean (SD) |  |
| Age | 37.37 (12.08) | 33.13 (13.00) | $t(93) = 1.647, p = 0.103$ |
| Years of education | 17.04 (2.47) | 16.68 (2.85) | $t(93) = 0.661, p = 0.510$ |
| Handedness (R / L / Mixed) | 45 / 4 | 44 / 1 / 1 | $\chi^2 = 1.644, p = 0.200$ |
| Gender (F / M) | 28 / 21 | 26 / 20 | $\chi^2 = 0.004, p = 0.951$ |
| FFMQ | 154.63 (15.48) | 134.11 (13.96) | $t(93) = 6.772, p < 0.001$ |

R: right-handed, L: left-handed, Mixed: ambidextrous, F: female, M: male, FFMQ: five-facet mindfulness questionnaire.

### 2.2 Procedure

To aid in understanding our overall procedure, a high-level overview is provided in Figure 1. EEG data were acquired via NeuroScan Acquire software and a SynAmps 2 amplifier, using a Neuroscan 64-channel Ag/AgCl Quick-Cap (Compumedics, Melbourne, Australia). Data were referenced online to an electrode between Cz and CPz. Electrode impedances were kept below 5 kΩ. The EEG was recorded from 60 electrodes (excluding CB1, CB2, M1, and M2) using the standardized 10/20 system at a sampling rate of 1000 Hz, with an online bandpass filter of 0.05 to 200 Hz. Data were collected across three minutes where participants were instructed to “rest, not meditate” with their eyes closed, and to let their mind do whatever it wanted, with no deliberate control exerted. All participants also completed five cognitive tasks within their EEG session (listed in the supplementary materials), which typically lasted between 2.5 and 3.5 hours in total. In the first study, resting-state data were recorded after the second cognitive task. In the second study, resting-state data was taken after the first task.

**Figure 1.**
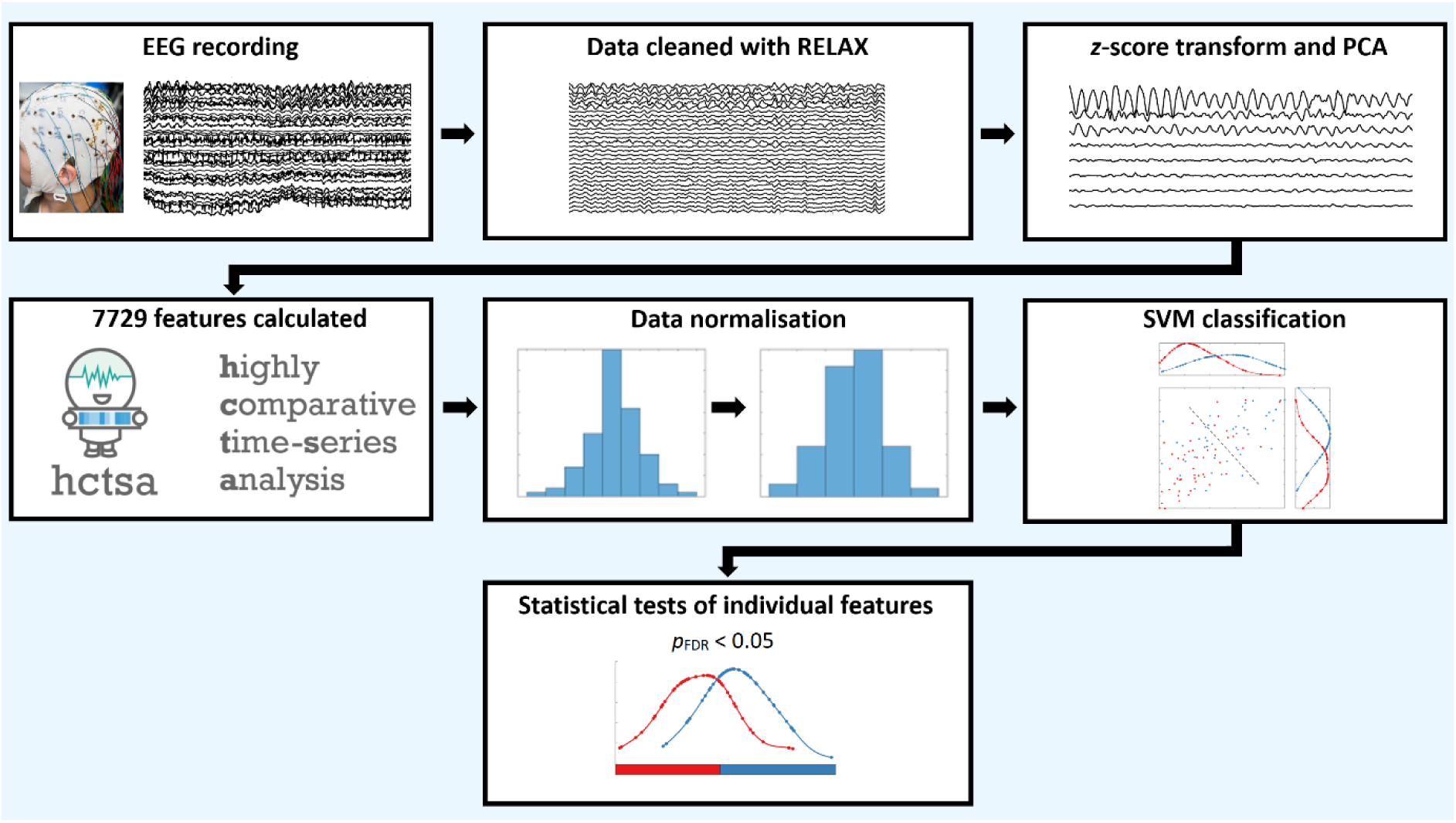
Depiction of the steps undertaken to pre-process the EEG data with the Reduction of Electrophysiological Artifacts (RELAX) toolbox, compute the highly comparative time-series analysis (*hctsa*) features, run the support vector machine (SVM) classification algorithm and test which features showed significant differences between the meditators and non-meditators. Note that *hctsa* feature extraction and classification was performed only on the top eight principal components. Between group differences in individual features were tested using Mann–Whitney U test statistics (Mann & Whitney, 1947). We controlled the false discovery rate to control for multiple comparisons using the method of Benjamini and Hochberg (1995).

After the EEG was recorded, a series of evidence-based steps was undertaken to optimally process the data based on previous research (Bailey et al., 2023c; Bailey et al., 2023f; Bailey et al., 2023d), which are reported in full in the supplementary materials. These pre-processing steps constituted: 1) automatic removal of all artifacts from the data while preserving the neural signal (Bailey et al., 2022a; Bailey et al., 2022b); 2) segmenting the data into 30 second epochs and automatically rejecting epochs showing any remaining artifacts (one non-meditator participant’s data was excluded at this stage as no artifact free 30s epoch could be obtained) (Bailey et al., 2023f; Decat et al., 2022); 3) baseline correcting the data (so the mean of the epoched data for each individual was zero); 4) downsampling the data to 160 Hz (which ensured the time-series features were calculated on the frequencies containing the majority of meaningful EEG variance); and 5) *z*-score transforming the data across all values within each epoch (which preserved the relationships between electrodes while normalising the amplitudes of the data so variations in amplitude did not bias the next steps).

While the feature-based classification approach requires a single (univariate) time-series from each participant to extract features, EEG data are typically recorded from many electrodes simultaneously (our EEG data were recorded from 60 electrodes). To address this, we followed a procedure established in our previous research where we used the dimensionality-reduction technique of principal component analysis (PCA) to extract highly explanatory time-series from spatially weighted principal components (PCs) (Bailey et al., 2023e). Within our dataset the top eight PCs explained 95% of the variance. Although PCA decomposition prior to the application of a machine learning algorithm is often performed after exclusion of the test set data to ensure the test set data is unseen by the PCA decomposition as well as the machine learning analysis, our primary aim was to use between group comparisons of individual features to determine which features provided the strongest differentiation between meditators and non-meditators. As such, the inclusion of all participants in the PCA decomposition ensured the individual statistical feature tests would be performed on PC weightings that were identical for all participants (in contrast to approaches that perform the PCA on each participant separately). Our approach also ensured the PCA decomposition was representative of activity obtained from all participants. However, to ensure that the inclusion of the test set data in the PCA did not bias our machine learning results, we performed 10 additional PCA decompositions, holding approximately 10% of the participants out from each decomposition (selected at random, analogous to a cross-validation test). The PC weightings produced by each of these 10 decompositions were essentially identical (the correlation between the weightings of the 10 PCA decompositions were all r > 0.999, see supplementary materials Figure S2), indicating that the inclusion of all participants in the PCA decomposition was unlikely to have biased our machine learning results or contributed to overfitting. Additionally, we performed an exploratory analysis of a representational set of the individual features that our primary tests showed to significantly differentiate the two groups, in which any PC that provided above chance classification accuracy was tested again using a cross-validation approach whereby the PCA decomposition excluded test set data. In this cross-validation approach, PC weightings were separately applied from each of the PCA training sets only to individuals from the test set in order to obtain the PC time-series, ensuring the test data were unseen by the PCA before the *hctsa* feature computation. Our machine learning and statistical tests of these features replicated our primary analyses, demonstrating that the PCA decomposition that included all participants did not bias our results or contribute to overfitting (these results are reported in the supplementary materials). Following pre-processing, we performed hctsa feature extraction, then fitted and evaluated the classification models.

### 2.3 *hctsa* feature extraction and normalisation

Version 1.07 of *hctsa*, which includes implementations of 7729 time-series features, was used with Matlab 2022b to extract features, then fit and evaluate classification models. *hctsa* performs different calculations to obtain outcome values for each of the >7000 features using Matlab functions and custom scripts provided within the *hctsa* software, with the full implementation and list of features provided in the software (Fulcher et al., 2023). *hctsa* includes implementations of statistical learning algorithms (including correction for multiple hypothesis testing) that can highlight the types of analysis methods that are best suited to classifying labelled classes of time-series data like EEG (Fulcher & Jones, 2017; Fulcher et al., 2013). Each feature was evaluated for each participant within each of the eight PCs to produce a single value for each feature within each PC and participant. A single value for each participant within each of the eight PCs was produced for each feature. Any non-real values or errors that were returned from the feature extraction were excluded from further analyses so that only valid data were included in our analyses. A feature was entirely excluded if it produced non-real or erroneous outputs across all participants. For example, within PC3, this resulted in the removal of 348 features, with 7381 features remaining for subsequent analysis (represented as a 95 x 7381 feature matrix). A similar number of features were removed within the other PCs. The *hctsa* feature matrix for each PC was normalized across all participants for each feature separately using a mixed sigmoid transform to enable more straightforward comparison of features measured on different scales and with different distributions (see supplementary materials for full details) (Fulcher et al., 2013).

After normalization, we used a linear support-vector machine (SVM) to classify meditators and non-meditators based on the >7000 time-series features for a given PC of the EEG data. To ensure that optimistic results could not be obtained from over-fitting, we used 10-fold cross-validation to calculate a mean balanced accuracy score. To control for class imbalance (our sample contains 49 meditators and 46 non-meditators) we used inverse probability class reweighting when training the SVM. To determine whether our classification accuracy was statistically significant, we used a permutation test with 1000 null samples. Each null sample involved shuffling the meditator/non-meditator labels across participants and obtaining a null balanced accuracy score from the same cross-validation procedure as used for the real labels (Henderson & Fulcher, 2022). The resulting null distribution allowed us to estimate a *p*-value as the proportion of null prediction accuracies that exceeded the real prediction accuracy. We used the false discovery rate (FDR) to control for multiple comparisons across the models obtained from the eight PCA components (Benjamini & Hochberg, 1995), and report FDR-corrected *p*-values as *p*_FDR_

### 2.4 Individual Feature Analysis and Feature Interpretation

After we determined which (if any) PCs were able to accurately classify the two groups, we next aimed to understand the types of dynamics that distinguish the meditators from non-meditators. While the multivariate classifier described above used normalized data from each of the >7000 features as the input, for the individual feature analysis, we analysed raw (non-normalised) individual feature values to interpret which individual features differentiated the two groups. This enabled clearer interpretation of the characteristics of each group for these features. To assess between group differences in individual features, we calculated Mann–Whitney U test statistics (Mann & Whitney, 1947) separately for each of the >7000 time-series feature from principal components that showed statistically significant classification accuracy. This allowed us to interpret which types of dynamical changes contained within the PCA component provided the strongest differentiation of the meditator and non-meditator groups. We controlled the FDR to control for multiple comparisons within any significant PC across all >7000 individual features within a PC using the method of Benjamini and Hochberg (1995). Given the large correlation between many features (Fulcher et al., 2013), we note that this approach (which assumes independence) is expected to yield conservative significance estimates.

Finally, to compare the performance of the *hctsa* features at distinguishing between the meditator and non-meditator groups to the performance of traditional canonical frequency band-power features, we tested the 10-fold cross-validation classification accuracy of each frequency band (delta: 1-4 Hz, theta: 4-8 Hz, alpha: 8-13 Hz, beta: 14-25 Hz and gamma: 25-45 Hz). To do this, we repeated the classification analysis described above, but instead of using the *hctsa* feature set, we used the averaged power within the 5 canonical frequencies as features. Firstly, we computed spectral power from each of the eight PCs separately using a Welch’s method, then averaged the power within each frequency band for each PC separately. We then included the averaged power from all five bands and all eight PCs in the classifier, which was represented in a 95 participant x 40 PC band-power matrix. Note that due to the smaller number of features in the band-power analysis, we included frequency measures from all eight PCs in the single classifier (in contrast to our primary tests of all *hctsa* features, in which we tested each PC separately to avoid potential overfitting issues caused by the inclusion of so many features in the classifier). Additionally, to match the analyses we conducted using the *hctsa* feature set, we also tested classifiers from the canonical frequency bands from each PC separately. Finally, we tested the exact Mann–Whitney U test on each individual PC band power and used the false discovery rate (FDR) to control for multiple comparisons across these 40 features using the method of Benjamini and Hochberg (1995).

Furthermore, because previous research has not typically applied PCA prior to frequency band analyses, we performed a frequency band analysis focused on specific electrodes. Unfortunately, doing so poses an issue of which electrodes to select (an issue that we addressed in our main analysis through the use of PCA dimension reduction). To address this issue, we selected single electrodes that either were reported by previous research to show maximal power within each frequency of interest, or were reported by previous research to show differences between meditators and non-meditators (delta: Knyazev, 2012, theta: Cohen and Donner, 2013, alpha: Lozano-Soldevilla, 2018, beta: Barry and De Blasio, 2017, gamma: Cahn et al, 2017). As such, we examined delta and theta power from FCz, alpha and gamma power from PO7 and PO8, and beta power from Pz.

## 3. Results

### 3.1 Cross-Validation Tests

The classifier trained on the 7381 hctsa features from the PC3 time-series showed above chance classification accuracy of meditators and non-meditators, which exhibited a mean balanced accuracy of 67% (SD across the 10 test folds = 6.54%, p_FDR_ = 0.007). None of the other classifiers trained on data from the other seven PCs showed statistically significant classification accuracy (all p_FDR_ > 0.10). The map of the spatial weightings and averaged power spectrum of the PC3 component across all participants is plotted in Figure 2. The power spectrum indicates a dominant alpha rhythm - the power spectrum averaged across all participants showed a peak frequency of 9.61 Hz. Peak frequency detected for each individual separately did not differ between groups (meditator mean peak frequency = 8.90Hz, non-meditator mean peak frequency = 8.64Hz, Mann-Whitney U p = 0.646). This is of interest, as alpha activity has been reported to be altered by meditation in previous research (Kerr et al., 2011; Wang et al., 2020). However, it is worth noting that the other seven PCs also showed a peak frequency in the alpha range (see supplementary materials Figure S1). The topographical map of PC3 showed that the electrodes that contributed the strongest weightings to PC3 were focused on midline central parietal electrodes, with a maximum at CPz (see Figure 2). Fronto-polar and temporal electrodes (where blink and muscle artifacts are respectively most prominent) did not provide strong contributions to PC3. As such, both the alpha prominence of the frequency spectrum and the topographical weightings suggest our classification accuracy was not influenced by blink or muscle artifacts, suggesting that the time-series features differentiating our groups were likely related to brain activity rather than non-neural artifacts.

**Figure 2.**
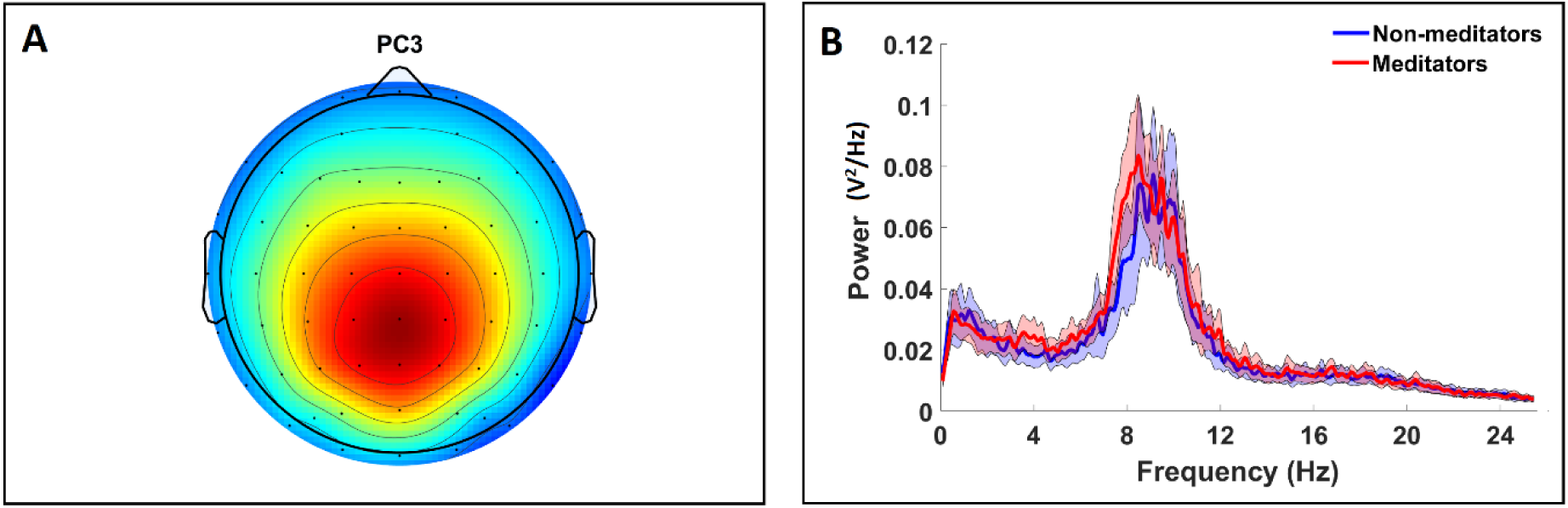
The topography and power spectrum for PC3. *Left*: The topographical map of weightings from each electrode that contributed to PC3. *Right*: the mean power spectrum for PC3 averaged across all participants within each group separately (computed using Welch’s method); shading reflects 95% confidence intervals.

### 3.2 Which time-series features differentiate meditators and non-meditators?

Within our Mann-Whitney U analyses of individual features, a total of 405 individual time-series features from PC3 showed significant differences between the meditators and non-meditators after controlling for 7381 multiple comparisons (*p*_FDR_ < 0.05). For visualisation and interpretation, we focused on the most discriminative 50 features (with *p*_FDR_ < 0.016), as these 50 features showed the strongest effects and captured the relevant types of methods contained in the full set of 405 significant features. Focusing on the top 50 features in this way allowed us to interpret the clusters of features that showed the strongest effects at differentiating meditators and non-meditators. To determine which types of time-series properties these features measured, we organised them using linkage clustering based on absolute Spearman correlations (|ρ|) between the values provided for each individual feature, implementing a cluster threshold of |ρ| > 0.75, which yielded groups of features that showed similar behaviour within our dataset. The relationships between these features and their interpretations are summarized in a dendrogram and cluster plot in Figure 3, which depicts several clusters of highly correlated features. Overall, there was a high level of correlation between different features, indicating that many features showed similar outputs to other features across participants within our dataset. The dendrogram (Figure 3) shows three large clusters of features that strongly distinguished the groups. Since these three clusters contained features that performed best at distinguishing the groups, and contained seven or more related features providing high performance, we focused our interpretation on these clusters. To provide a parsimonious interpretation of these features, we selected representative and clearly interpretable exemplars for each of these three clusters to explore in more detail. We note that the exemplars we selected also provided the strongest statistically significant differences between the two groups within each respective cluster. However, we note that there were features within the 405 significant features that did not correlate highly with the three clusters we interpret, and these features may provide further novel insights into differences in brain activity between meditators and non-meditators. We discuss some of these features briefly in our supplementary materials, and provide a full list of the significant features and their correlation matrix in our supplementary materials.

**Figure 3.**
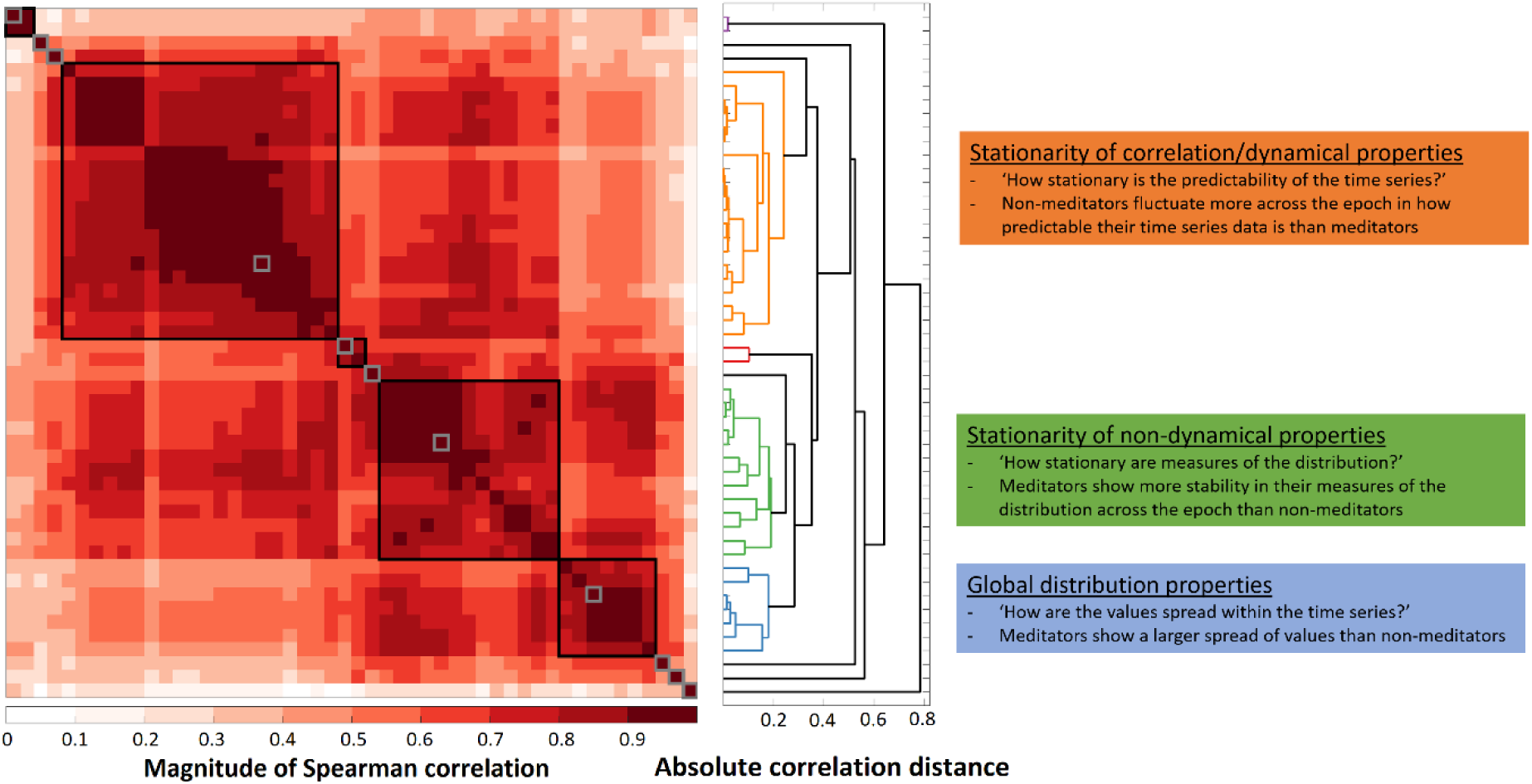
*Left*: the cluster plot of the top 50 features from PC3, clustered using absolute Spearman correlations and a threshold of |ρ| = 0.75 for forming clusters, enabling visualisation of how strongly the different clusters are related to each other. *Middle*: dendrogram of the top 50 features, clustered using absolute Spearman correlations and a threshold of |ρ| = 0.75 for forming clusters. This plot groups features by the strength of the correlation with other features, such that those with smaller absolute correlation distances are depicted in the same colour, and these features show a strong linear relationship between the values from each feature. *Right*: Annotated brief interpretations are provided for the three largest clusters. The full list of significant features from PC3 and correlation matrix can be found in the supplementary materials.

The first cluster of features from PC3 (coloured orange in Figure 3) contains features that assess the stationarity in the dynamic properties of the time series, in particular, the consistency of time-order-dependent statistical patterns across shorter subsegments of the full 30s recording. These features implement measurements derived from the concept of ‘stationarity’ in time-series analysis, which can loosely be thought of as the extent to which the statistical properties of the time series are constant. Practically, for finite-length real-world time series, this often involves quantifying the consistency of statistical properties measured in local subsegments of a recording. The 17 features from this cluster indicate that meditators displayed more stable and consistent dynamical patterns within PC3 across the 30s EEG epoch than non-meditators. An example feature in this cluster measured the variability in the median-based time-series predictability across five non-overlapping (6 s-long) segments of the 30s time series. This feature is named **FC_LocalSimple_median7_sws** in *hctsa*. We refer to it in this article as the ‘stationarity of median-model predictability feature’ for simplicity. This feature assesses predictability by computing the median of 7 consecutive samples, then using that median to forecast the next time-series sample. Comparing the standard deviation of the residuals from this prediction process across five non-overlapping (6s) segments of the time-series, meditators showed a more consistent level of residual variance across segments than non-meditators (Figure 4A, and a schematic explanation of the computation of this feature is provide in the Supplementary Materials Figure S3). This feature displayed the strongest ability out of all *hctsa* features at distinguishing meditators from non-meditators (*p*_FDR_ = 0.012, Cohen’s *d* = 0.911, Figure 4A). The full list of significant features is provided in the supplementary materials, and readers can refer to *hctsa* for details of how each individual feature was computed (Fulcher & Jones, 2017). Other types of conceptually similar features in this cluster of features (orange in Figure 3) also measured the consistency of different types of dynamical properties across shorter segments of the time series and yielded similar insights. Overall, these features indicated that the meditator group showed more consistent dynamical properties within the PC3 component of their EEG signal than non-meditators across this 30s timescale, where the time-order dependent statistical properties of the time-series were less variable between the different 6s segments of the data.

**Figure 4.**
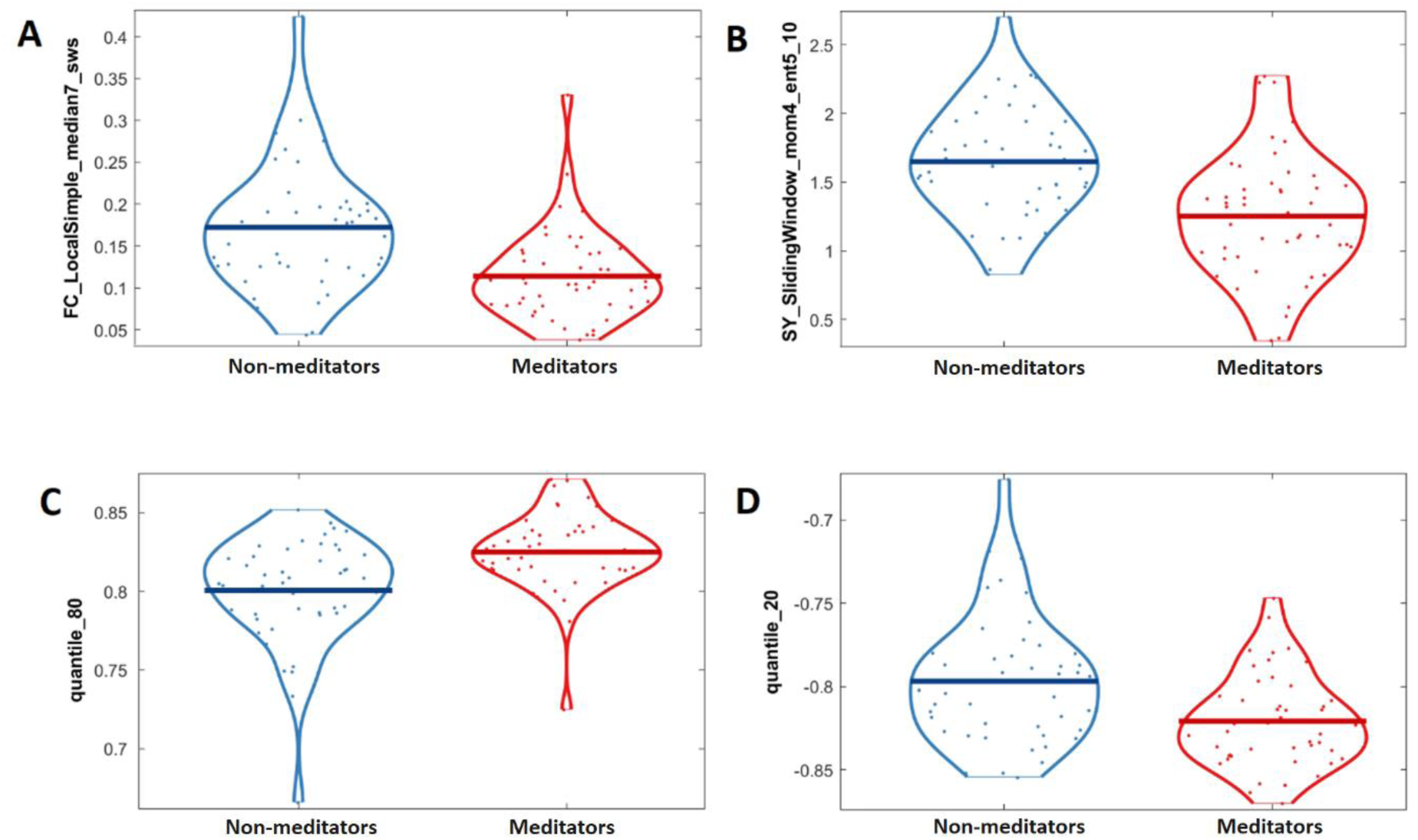
The distribution of individual feature values obtained from PC3 for the meditator and non-meditator groups for top-performing features reflecting the three clusters depicted in Figure 3 shown as violin plots. A: the **FC_LocalSimple_median7_sws** feature (or ‘stationarity of median-model predictability’). This feature reflects the consistency in a simple median-model predictability of participant’s PC3 time-series dynamics, so is thus a measure of stationarity in the dynamic properties of the time-series, with lower values reflecting higher stationarity. Meditators showed higher stationarity in how predictable their PC3 data were between each six second segment (p_FDR_ = 0.012, Cohen’s d = 0.911). B: The distribution of values from each group for the **SY_SlidingWindow_mom4_ent5_10** feature (or ‘kurtosis stationarity’). This feature assessed how variable the kurtosis of time-series values was across overlapping 6s segments of the epoch within the PC3 time-series, with lower values reflecting higher stationarity. Meditators showed lower values for this feature, indicating more consistency (or stability) in the distribution of their data (p_FDR_ = 0.012, Cohen’s d = 0.956). *C*. The distribution of values from each group for the **quantile_80** feature. This simple feature provided the value of PC3 at the 80th percentile of the *z*-scored time-series. Meditators showed higher values for this feature (*p*_FDR_ = 0.012, Cohen’s d = 0.842). *D*. The distribution of values from each group for the **quantile_20** feature. This simple feature provided the value of PC3 at the 20th percentile of the *z*-scored time-series. Meditators showed lower values for this feature (*p*_FDR_ = 0.032, Cohen’s *d* = 0.658).

The second cluster of features from PC3 that performed well at differentiating meditators and non-meditators (coloured green in Figure 3) were also related to the consistency of the statistical characteristics of the time-series across different segments of the data (providing another measure of stationarity). However, the features in this cluster mostly captured ‘non-dynamic’ properties of the data, which depend on the distribution of values in a time-series segment, rather than their temporal ordering (where the distribution is the shape created by plotting each value from the 30s epoch of PC3 on the x-axis and the frequency of the occurrence of that value on the y-axis). Our results showed that the meditation group had more consistency in their distribution of time-series values across different segments of PC3 compared to non-meditators, effectively showing higher stationarity in the distributional properties of their time-series. An example feature within this cluster, with the largest effect size in this cluster, is named **SY_SlidingWindow_mom4_ent5_10** in *hctsa*, and is referred to as ‘kurtosis stationarity’ for simplicity here. This ‘kurtosis stationarity’ feature measured the variability in the kurtosis of time-series values across overlapping 6s segments of the epoch (where 90%-overlapping sliding windows were used to obtain the data segments, and variability was quantified using kernel-smoothed entropy). Kurtosis loosely measures the shape of a distribution’s tails and peak (i.e., how ’peaked’ or ‘flat’ it is) relative to a Normal (Gaussian) distribution (DeCarlo, 1997). Positive values for kurtosis are obtained when distributions are more peaked (with more values centred around the mean) than a Normal distribution, and negative kurtosis values are obtained when distributions are less peaked (‘flatter’) and have more values in the tails of the distribution away from the mean. Our results showed that meditators had more consistent kurtosis estimates within PC3 across local (6s) time-series windows than non-meditators (*p*_FDR_ = 0.012, Cohen’s *d* = 0.956, see Figure 4B).

To aid with a practical understanding of this difference, distributions of PC3 values from five non-overlapping 6s segments from four exemplar participants are plotted in Figure 5. Inspection of these distributions shows that participants with higher values for this ‘kurtosis stationarity’ feature showed more variability in distribution shape across the 6s segments than those who showed lower values for this feature. Most of the remaining features in this second cluster (green in Figure 3) yielded qualitatively similar insights, capturing the consistency of local kurtosis measurements in slightly different ways (including variations in how the segments of the data were selected, and the use of standard deviation instead of entropy to quantify variability across windows).

**Figure 5.**
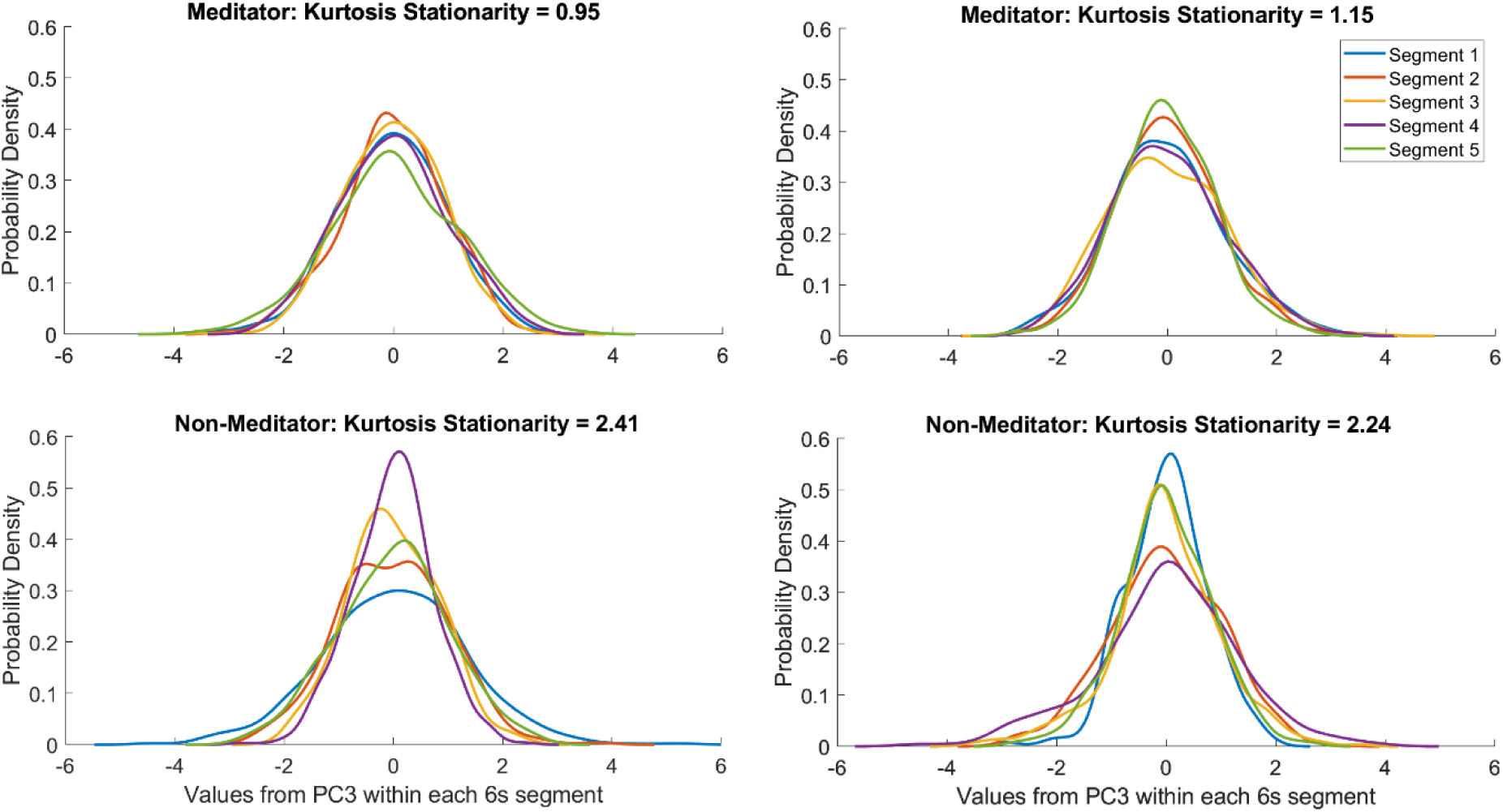
Distributions of PC3 values from five non-overlapping 6s segments from four exemplar participants demonstrating high and low ‘kurtosis stationarity’ values, with the ‘kurtosis stationarity’ (**SY_SlidingWindow_mom4_ent5_10**) values obtained from these four participants provided in the title for each plot. Note the larger variability in distribution shape from the 6s segments for participants that showed higher values for this ‘kurtosis stationarity’ feature, providing a visual depiction of how variability in the distributional shape across different 6s segments affects the ‘kurtosis stationary’ feature.

Since PC3 contained a prominent peak of oscillatory power in the alpha band, we conducted a brief exploratory analysis of both the ‘stationarity of median-model predictability’ and ‘kurtosis stationarity’ features after data were bandpass filtered to remove all frequencies outside of the alpha band. This demonstrated that both features were highly related to stationarity in alpha oscillations (reported in full in the supplementary materials). The ‘stationarity of median-model predictability’ feature showing almost identical results after data were bandpass filtered to only contain alpha activity, while the ‘kurtosis stationarity’ feature showed a similar effect size but a lower correlation between the alpha-filtered and non-filtered data. This result suggests that the higher stationarity in the meditator group might be driven by higher stationarity of their alpha oscillations.

The third cluster of high-performing features from PC3 (coloured blue in Figure 3) measured properties of the shape of the distribution of *z*-scored time-series values (from the entire 30s epoch). These features indicated that the meditator group were more likely to display PC3 time-series values with a distribution where the 20% to 3% most extreme values deviate further from the mean than non-meditators. This is consistent with a pattern of activity from the PC3 component of EEG data that contains larger (or longer) voltage peaks and troughs. The feature in this cluster with the largest effect size (labelled **quantile_80** in *hctsa*, *p*_FDR_ = 0.012, Cohen’s *d* = 0.842) measured the 80th percentile value of the *z*-scored time series (i.e., after ranking all the values of the *z*-scored time series by magnitude). As such, **quantile_80** is a measure of distribution shape, capturing (in units of standard deviation) how far the 80th percentile of values from the time series are from the mean (where the mean is 0 for the *z*-scored time series). Meditators also showed significantly lower values than non-meditators for the 20th percentile (named **quantile_20** in *hctsa*, *p*_FDR_ = 0.032, Cohen’s *d* = 0.658, Figure 5D), although with a smaller effect size than **quantile_80**. As shown in Figure 4C and Figure S4, relative to their mean and standard deviation, the 20th and 80th percentile of deviations in the PC3 signal were further from the mean for the meditation group than for non-meditator group. However, it is worth noting that the non-meditator group showed non-significantly higher values for the quantile_1 and quantile_99 features than the meditator group, indicating that meditators did not show more extreme values in the 1% most extreme values from the mean. Other features in the third cluster (labelled blue in Figure 3) represented conceptually similar global measures of distributional shape, including measures of histogram entropy and outliers. As such, these other features also captured differences in the distribution shape of the overall values in the PC3 time-series data between meditators than non-meditators.

While these three clusters assessed conceptually different characteristics of the data, it is also worth noting that the features from the different clusters were often also strongly correlated. This was particularly the case for the second and third clusters, where the correlation between features in the cluster that included measures of the stationarity of distributional properties of the data and features in the cluster that included measures of the distributional properties of the entire time-series was often ρ > 0.5, for example the correlation between **quantile_80** and ‘kurtosis stationarity’ features was ρ = 0.632 (p < 0.001).

### 3.3 Comparison to features derived from power in canonical frequency bands

Finally, to determine whether traditional frequency band power could provide a similar differentiation of the two groups to the top performing features from the *hctsa* feature list, we tested the performance of five canonical frequency band-power features (delta, theta, alpha, beta and gamma) computed from PCs 1-8 at classifying meditators and non-meditators. Our analysis of the canonical frequency band power across all PCs together in the same classifier provided a balanced accuracy of 56% (SD = 12.20%, *p* = 0.332). Further, when classification models were trained on the canonical frequency band power from each of the PCs separately, in contrast to the results of our analysis with *hctsa*, none of these classifiers from individual PCs provided significant balanced classification accuracy (all *p*_FDR_ > 0.05). When examining power averaged within individual canonical frequency bands (i.e., delta, theta, alpha, beta and gamma bands) from individual PCs, only the alpha band for PC6 passed an uncorrected significance threshold (*p* = 0.0079, reported uncorrected here to demonstrate the replication of previous research). None of the values from any PC within the canonical frequency bands significantly differentiated the two groups after controlling for multiple comparisons across the 40 comparisons (all *p*_FDR_ > 0.3). Similarly, the analysis of band power measures extracted from single electrodes did not show significant classification accuracy (balanced accuracy = 59.91%, *p* = 0.080). Analysis of the different band power measures showed that no band power measure significantly differentiated the two groups (all *p*_FDR_ > 0.10). However, prior to correction for multiple comparisons, alpha and beta frequency power showed significant differences, with meditators showing higher frequency power for both frequencies (*p* = 0.0417 and *p* = 0.0323 respectively).

It is worth noting that the distribution and stationarity features from *hctsa* provided results that were significant above the much higher multiple comparison control threshold across all *hctsa* features (which included an order of magnitude more individual features than the number of oscillation band features). These results indicate that the novel distributional and stationarity-based features of the EEG data (uncovered by *hctsa*) provided a stronger differentiation of meditators from non-meditators than the canonical frequency band power analyses.

## 4. Discussion

Our results showed that an individual could be classified as an experienced meditator or non-meditator with above chance accuracy (67%, *p*_FDR_ = 0.007) using one component of a single 30s epoch of EEG activity (with a central-parietal maximum) and a comprehensive feature-based representation of the time-series. For this component of the EEG data, a total of 405 individual features significantly differentiated the two groups, even after stringently controlling for multiple comparisons across >7000 candidate features. Our exploration of these individual features showed that the largest differences between the groups were related to differences in the stationarity of the time-series (both of dynamical properties and distributional shape) and features related to the global distributional shape (particularly reflecting how values were concentrated towards the mean compared to how they were concentrated towards the extremities). These results provide further support for the considerable amount of research demonstrating that a meditation practice sustained over years is associated with alterations to brain activity (Cahn & Polich, 2006; Falcone & Jerram, 2018; Ganesan et al., 2022).

However, as far as we are aware, most of the features uncovered by our systematic, data-driven approach have not previously been used in meditation research and they have only minimally been explored in EEG research more broadly. Furthermore, conventional EEG analysis methods based on the linear correlation structure (using classical frequency-band power features) were not amongst the significant features, and if we had analysed our data using band-power measures we would have concluded that the groups did not show different brain activity. Indeed, any features derived from a Fourier transform assume stationarity, and are thus insensitive to differences like the distinctively higher stationarity that characterized the PC3 component of the meditation group’s EEG signal. Our results thus demonstrate the usefulness of characterizing EEG time series using methods that go beyond methods that assume an underlying stationary process that is well-characterised by its Fourier power spectrum. In contrast to these conventional EEG methods, our top-performing features captured novel types of time-series properties that may provide a new lens through which to understand the effects of meditation, as well as suggesting promising novel application in understanding brain states in general. As such, our analysis provides important new insights that could be productively used to explore the mechanisms by which mindfulness practice affects mental health and cognition. Additionally, we are not aware of any studies within the field of EEG research that have used measures of within epoch univariate time-series stationarity to classify groups, or test for differences between groups; therefore, our results highlight a valuable new direction for EEG research more generally. In particular, the predominant use of band-power features to represent EEG dynamics may be a factor in studies presenting null results, with recent examples including null results for the classification of individuals with Autism from neurotypical individuals using a very large dataset (Dede et al., 2023). In contrast to these null results, our ability to significantly classify meditators is the result of our use of an expanded methodological toolbox (provided by *hctsa*) — other studies may also benefit from leveraging more comprehensive statistical summaries of neural dynamics.

Our analysis of the individual features from the third principal component of the EEG data that differentiated meditators and non-meditators showed one high-performing cluster of features which assessed the stationarity of the dynamical properties of the time-series across shorter local segments of the 30s epoch. This cluster of features revealed that meditators have more consistently predictable time-series within the PC3 component of an EEG epoch – essentially, within the meditator’s PC3 component, the time-order dependent patterns within the data were more similar across different 6s segments of the full 30s epoch compared to non-meditators. Features assessing the consistency of distributional properties of the time-series also strongly differentiated the two groups, with meditators showing more consistency in the distribution of values during different segments of the PC3 component. Our ability to contextualise this finding is limited, as we are not aware of any other research that has directly assessed the temporal stability of EEG data for comparison with our results. However, one potential implication of the higher level of stationarity may be that meditators generate more temporally stable neural activity within the brain regions represented by the PC3 component (e.g., across the 6 second segments of the 30 second EEG recording, as was assessed by many of the features in our study). This higher degree of temporal stability might be a biological marker of neural activity that underpins the enhanced stability of attentional focus suggested to arise from meditation practice. Further research is required to explore this possibility.

Features that measured how the data were globally distributed also significantly differentiated meditators and non-meditators, with meditators showing a broader distributional shape of PC3 time-series values, or higher densities of values further away from the mean. This result is consistent with larger, longer or more frequent non-outlying voltage peaks and troughs generated in the brain regions that contributed to the third principal component of their EEG activity. While we are not aware of EEG research that has examined distributional features, research applying *hctsa* to examine resting-state functional magnetic resonance imaging (fMRI) time-series has detected effects in distributional features similar to those detected in our study (Shafiei et al., 2020). Shafiei et al. (2020) showed that the variation of fMRI time-series features related to distributional entropy and kurtosis (akin to a group of features that distinguish the meditation group) explained a large amount of the variability in the anatomical properties of the brain. The spatial variation of these distributional shape-based properties of the fMRI signal was correlated to similar spatial variations in cortical thickness, intracortical myelination, and connectivity hierarchies within brain networks (Shafiei et al., 2020), flagging the distributional shape-based properties as an underappreciated signature that mirrors underlying anatomical variation. It is worth noting that fMRI data is typically recorded at a resolution of 1-3 samples every second (in contrast to 160 samples per second as per our results), and it is not clear how haemodynamic and electrophysiological signals are related (although some work has shown correspondences between correlation timescales estimated from magnetoencephalography and fMRI; Watanabe et al. (2019)). Nonetheless, both our study and the study by Shafiei et al. (2020) highlight the relevance of quantifying the shape of the distribution of fluctuations in resting-state neural activity—a property that is simple to interpret but remains underexplored.

It is also worth highlighting that the topographical map of the PC3 component showed a maximum in midline central-parietal electrodes. PC3’s topographical map is almost identical to the topographical pattern shown by “microstate E”, within research into EEG microstates (Tarailis et al., 2023). Microstate E has been reported to be generated by activity from the bilateral medial prefrontal cortex, including the dorsal anterior cingulate cortex, superior frontal gyrus, bilateral middle prefrontal cortex and bilateral insula (Tarailis et al., 2023). Both microstate E and the brain regions underpinning its generation are thought to comprise part of the salience network, which is associated with interoceptive and emotional processing (Tarailis et al., 2023). Interception and emotional processing have been suggested to be enhanced by mindfulness meditation practice (Brewer & Garrison, 2014; Garrison et al., 2015; Tang et al., 2015). As such, our result indicating that meditators show more temporally stable activity and longer, larger, or more frequent non-outlying voltage deviations in PC3 may suggest meditators can be characterised by more short-term temporal stability and a larger range of activity in brain regions underlying interception and emotional processing.

Given the novelty of our results with regards to the univariate stationarity and distributional properties of the data, there are no directly comparable studies within the meditation literature to help guide our interpretation. However, previous research has reported that experienced meditators in the meditation state show increased stability in the topographical pattern of EEG activity and in the temporal patterns of both EEG and fMRI based connectivity (Escrichs et al., 2019; Toutain et al., 2020; Vivot et al., 2020). The increased fMRI connectivity stability finding has also been reported in novice meditators, and for comparisons between meditator and non-meditator resting-state connectivity (Escrichs et al., 2019). Previous research has also shown that meditators generate more consistent brain activity in response to stimuli while performing a cognitive task, enabling their brain activity to time-lock more effectively to the stimuli presented in the task (Bailey et al., 2023b; Lutz et al., 2009). However, it is important to note that these measures of topographical or connectivity-based stability may be unrelated to the features that differentiated meditators and non-meditators in our study.

The aforementioned studies also mostly examined neural stability during the meditation state compared to the resting state, rather than comparing resting-state activity between meditators to non-meditators. Despite these differences, the meditation-state related increases in topographical or connectivity related stability that they report provides a possible suggestion for the cause of the increased stability in meditator’s neural activity — it could be that prolonged practice of increased stability of neural activity during meditation leads to higher levels of stability in neural activity. For example, it may be that meditation increases the engagement of specific attention related brain regions, which, with repeated practice, might exert a regulatory effect on other neural processes, leading to a durable increase in neural stability. Alternatively, it may be that the increased integration of brain networks during meditation practice (van Lutterveld et al., 2017) provides enhanced regulation of brain activity from moment to moment, enabling increased stability of meditator’s neural activity. In support of this suggestion, a growing amount of recent research suggests that the short-term dynamics of neural connectivity are predictive of attention function, both within and across individuals (Liu et al., 2018; Song & Rosenberg, 2021).

An additional mechanism that may explain the increased stability of meditator’s neural activity has been suggested through modelling work reported by Saggar et al. (2015). Their work simulated EEG data based on a model of the interaction between cortical, corticothalamic and intrathalamic brain regions. Saggar et al. (2015) compared the effects of adjustments to these simulated interactions to the effects detected within real EEG data recorded before and after two separate groups undergoing 3-month-long meditation retreats. They found that changes in the EEG data from before to after the retreat were partially explained by a model that contained a decrease in inhibition of the secondary relay nuclei by the thalamic reticular nucleus. This decreased inhibition by the thalamic reticular nucleus provided increased dynamic stability of the modelled EEG activity. While modelling approaches are only approximate, can only model a limited resolution and limited number of brain regions, and rely on a number of assumptions, the thalamus *has* been suggested to be involved in attention regulation in meditation, acting as a filter for sensory inputs prior to their reaching the cortex (Saggar et al., 2015). Future research could explore a comprehensive explanation involving increased stability of thalamic gating providing increased focused attention towards sensory sensations and the increased stability of neural activity in meditators (within the variance represented by the PC3 component) found in the current study.

It is also worth noting that the stationarity measures from PC3 detected in our study were highly influenced by activity within the alpha oscillatory frequency band, with our exploratory analysis (reported in the supplementary materials) showing similar effects in the stationarity measures after data were bandpass filtered to exclude frequencies outside of the alpha frequency range. Previous research has suggested that alpha activity provides two functions. Firstly, it provides an active top-down inhibitory function, whereby higher-order (cognition-related) regions inhibit non-task related lower-order (sensory processing) regions. Secondly, alpha activity provides a timing function, whereby the phase of alpha oscillations is timed to enable specific neurons to fire (Klimesch, 2012). Both the inhibitory and timing functions modulate the ‘gain’ of neural signals from lower order brain regions, enabling attention enhancement (Klimesch, 2012). Recent research using intracranial electrodes has also shown that alpha oscillations propagate hierarchically from higher order regions to lower order regions across the cortex, propagating from parietal to occipital visual processing regions, from associative to primary regions within the somatosensory cortex, and from the cortex to the thalamus (Halgren et al., 2019). Within these regions, the combination of alpha activity’s inhibitory and timing functions may fulfil a top-down gating role for the propagation of sensory information from lower-order sensory regions to higher-order brain regions (Halgren et al., 2019). As such, one potential explanation for our results that would be worth testing in future research might be that because of their attention training, meditators demonstrate increased stability of the alpha oscillation mediated influence of higher order regions of the cortex to lower order regions. It is worth noting that all PCs showed prominent alpha peaks, so the putative increase in alpha stability may be spatially specific, only differentiating the groups based on the weightings of PC3. However, while only PC3 provided above chance classification of the two groups, our brief inspection of the features from the non-significant PCs indicated the stationarity and distributional features were amongst the top performing features for all PCs, with the same direction of effects (although no individual features from the other PCs passed our stringent multiple comparison control thresholds). This may suggest that while PC3 captured the most important variance in stationarity and distributional features of the data for differentiating meditators and non-meditators, the stationarity and distributional properties of neural activity in brain regions outside of the regions that provided strong weightings to PC3 may also differ in meditators (although with smaller effect sizes). Unfortunately, due to the diffuse nature of the neural activity measured by EEG recordings, we cannot confidently ascertain the brain regions likely to be generating the activity captured by PC3. Source localisation focused on the features we showed differentiated the two groups in PC3 may be able to determine whether our results are regionally specific and which regions might be responsible for our results.

Regardless of how the effects detected in our study arise, our findings related to stationarity suggest that it may be fruitful for large-scale mediation analyses to explore meditation’s neural mechanisms of action using measures of the stability of neural activity. If increased stability of neural activity does reflect a neural mechanism of action, then interventions could be designed to target this characteristic of neural activity, potentially leading to more effective interventions. Future research could also determine whether conditions that show lower stationarity in their EEG might be recommended to mindfulness practice as a method to resolve this potential pathophysiological marker. For example, lower temporal stability of EEG patterns has been associated with less self-control and higher risk-tasking behaviour (Kleinert et al., 2022). Future research could also explore whether the time-series features that capture different aspects of stability across different timescales of neural activity may also offer the potential to detect progression in a meditation practice.

While measures of distributional shape and stationarity best differentiated meditators and non-meditators, our results revealed significant group differences in a range of other time-series properties, and other features may be interesting to explore. In particular, we were interested in complexity or signal entropy measures, such as Approximate Entropy, Sample Entropy, Permutation Entropy, and Lempel–Ziv complexity. These measures have received increased attention in recent neuroscience research as measures of the uncertainty or information content within the EEG signal. Such methods have been proposed to relate to the richness of conscious awareness, with increases in conscious awareness suggested to be associated with greater signal complexity (Carhart-Harris, 2018) and decreases in these measures suggested to reflect a single pointed attentional focus (Young et al., 2021). Our analysis did not show significant group differences in these types of signal entropy measures (although non-dynamic entropy-related features which were used to quantify properties of the distributional shape did show differences between the groups, and entropy-related features used to quantify variation in predictability across windows as sliding-window stationarity did show differences between the groups). The lack of difference between meditators and non-meditators in these signal entropy measures is in contrast with some previous research, which has reported higher Sample Entropy or Lempel–Ziv complexity both during the meditation state and in long-term meditators compared to non-meditators or novice meditators (Lu & Rodriguez-Larios, 2022; Vivot et al., 2020). There are differences in data pre-processing between our research and previous research, with previous research examining data from electrodes rather than analysing principal components, which may explain this conflict with previous research. However, the entropy-related findings are not consistent in the literature, with some research reporting lower Lempel–Ziv complexity during the meditation state compared to the mind wandering state in experienced meditators (Young et al., 2021).

### 4.1 Limitations and future research

The primary limitation of the current study is that the data were collected using a cross-sectional design. This approach provided a benefit in that the meditators were more experienced than is feasible within longitudinal research, and thus likely showed more robust and larger differences in brain activity than an 8-week mindfulness intervention could elicit. However, the cross-sectional design precludes us from inferring causality from our findings. Further, only one EEG recording was analysed from each participant, and only the first 30s of artifact free EEG data were analysed, so we did not assess whether the measures we assessed were consistent across different days/weeks/months. However, this limitation is also a strength of the current study, in that it indicates the potential robustness of differences in neural activity in the meditation group – only 30 seconds of EEG data was sufficient to accurately classify meditators and non-meditators. Future research might benefit from the inclusion of multiple epochs, potentially across multiple days, to obtain more consistent measurement of the relevant features, assess how consistent these features are across time, and determine whether variability over time in the features might also differentiate the groups.

Additionally, our approach attempted to maximise both generalisability (across different meditation practices) and statistical power to detect meaningful effects (including participants from the broader category of mindfulness meditation allowed us to obtain a larger sample size). As such, we included all mindfulness meditation type practices, and did not separate practitioners by focused attention vs open monitoring practices, for example. In this regard, our approach contains both a strength – where we identified strong and generalisable differences in brain activity across all mindfulness meditation techniques, and a weakness – we could not separate differences in brain activity by the type of practice. However, we note that while it would be interesting to examine focused attention and open monitoring practices separately, the differences between the two practices are likely to be much smaller than the differences between meditators and non-meditators, as such, doing so would require much larger sample sizes to detect any effects. Additionally, our perspective is that there is considerable conceptual overlap between ‘focused attention’ and ‘open monitoring’ practices, as open monitoring requires training in focused attention to be useful, and because open monitoring could be considered as simply a focused attention practice with no specific object of focus. Other researchers have similarly noted conceptual ambiguity in delineating these supposedly separate meditation practices (Schoenberg & Vago, 2019). Similarly, we did not perform any analyses split by sex or gender, as our sample size was not sufficiently powered for this split analysis. As such, our results provide no indication of whether the same effects are present across different sexes or genders.

It is also worth noting that *hctsa* computes features from a single (univariate) time series. However, EEG data were recorded from 60 electrodes. To address this, we performed a PCA to reduce the data to eight components explaining over 95% of the variance in the data, drastically reducing the number of multiple comparisons relative to examining each electrode separately and allowing us to analyse spatially distributed and highly explanatory patterns that may not have been present in any individual electrode. To ensure a weighting consensus, the PCA decomposition was performed on a matrix that included data from all participants in a single matrix, ensuring that the PC weightings were common to all participants. This allowed us to take a comprehensive list of over 7000 features and narrow it down to provide a smaller but novel list of conceptually distinctive and informative features from a single principal component that are highly effective at detecting differences in brain activity between meditators and non-meditators. This highlights the value of our data-driven and hypothesis-free approach (further explanation of this point is provided in the supplementary materials). This approach is likely to be valuable to translate to other methodologies within neuroscience, where data often consist of many concurrent (multivariate) time-series, for example fMRI. The approach is also likely to be useful in addressing other problems, for example, detecting the mechanisms underlying both the cause of and treatments of major depression.

Despite the advantages of our approach, the relationship between activity at any electrode and the underlying sources of brain activity is not straightforward to interpret. As such, although the topographical map for PC3 showed a central-parietal activation that is similar to a microstate topography previously reported to be generated by the bilateral medial prefrontal cortex, we cannot be certain about which exact brain regions generated the activity. While source localisation has limitations, it could be used to estimate which brain regions might have been responsible for the differences we detected. Alternatively, magnetoencephalography might be a useful tool to obtain better spatial resolution. Additionally, while we examined eight univariate component time-series, the brain is an interconnected network, and consciousness, cognition, mental well-being, and the effects of meditation are related to these interactions between brain regions (Boly et al., 2012; Ganesan et al., 2022; Miljevic et al., 2023; Thivierge & Marcus, 2007). Research has demonstrated that meditation is associated with differences in the characteristics of the functional connectivity across the brain (Ganesan et al., 2022), and that attention is predicted by the dynamic modulation of connectivity within the brain (Liu et al., 2018; Song & Rosenberg, 2021). As such, it is likely to be valuable to implement an analogous highly comparative approach to comprehensively assess pairwise connectivity measures in future work (Cliff et al., 2022), or to use deep learning methods to extract differences in connectivity patterns between meditators and non-meditators, and to assess whether their combination with the univariate component features from the current study could yield further improvements in classification performance.

As with all research, replication of our results is necessary before the insights from our study could be applied in practice. To enable replication of our research, we have provided the full list of PC3 features that significantly differentiated the two groups after stringent multiple comparison controls in our supplementary materials section (along with how well these features correlated to one another). The code we used to clean the EEG data are publicly available (Bailey et al., 2022a; Bailey et al., 2022b), and we have provided the principal component weightings we used to transform our data in our supplementary materials, so if another dataset were collected, the same data transforms could be applied in order to replicate our approach.

We also note that alternative parameter settings for specific features may provide an even stronger ability to detect differences between the neural activity of meditators and non-meditators. Our brief exploration of alternative parameter settings (reported in the supplementary materials) suggested that the relevant timescale for the higher ‘kurtosis stationarity’ shown by the meditators involves data segments in the 2 to 6 second range. It may also be interesting to look at data epochs longer than 30 seconds, to determine how localised in time the stationarity effects are. More broadly, the detection of between-group differences in measures of the stationarity of the EEG signal suggests that it is likely to be promising for future to explore the variability of statistical properties of EEG data in other populations, and within other states (cognition-related states, different states of consciousness, and in pharmacologically induced states). While many traditional EEG analysis methods are derived from a Fourier power spectrum of a full recording, thus assuming stationarity of the dynamics on that timescale, our results indicate that meaningful effects can be detected from directly measuring stationarity using algorithms that assess the consistency of statistical properties across shorter windows of the recording. This is analogous to the trajectory of fMRI research methodology, which traditionally computed functional connectivity (examining correlations in activity between brain regions) using the entire recording (assuming that connectivity between regions was approximately stationary). More recent approaches have analysed the variability within these connectivity measures across shorter timescales to provide measures of ‘dynamic functional connectivity’, the results of which indicate that this timescale of stationarity/variability is informative of different states and conditions (Demirtaş et al., 2016; Hindriks et al., 2016; Jones et al., 2012; Matsui et al., 2022; Zalesky et al., 2014). Our results indicate that a similar conceptual framing of variability of statistical properties within windows of individual component time series can be similarly valuable.

Finally, the list of features we have provided could productively be used to explore the mechanisms by which mindfulness has beneficial effects on mental health and cognition. If these measures are identified as reflective of the mechanisms by which mindfulness meditation has beneficial effects, then these mechanisms could be used to test the design of optimally effective mindfulness interventions, as well as potentially assess meditation progress. Individuals who show impairments in these mechanisms might also be recommended to mindfulness interventions as a treatment to resolve the impairment. Further research into these mechanisms might also enhance our understanding of brain function in general. The measures of stationarity we have detected might also be promising targets for neurofeedback (an example of how this might be applied is provided in the supplementary materials). If this neurofeedback approach is successful, it might be associated with similar well-being and attention benefits to those reported by and detected in experienced meditators. However, it is also possible that ‘device-oriented’ approaches like this may be a distraction from a meditation practice that may already be optimal when taught within certain traditions, so it may be that this proposed neurofeedback approach would be more suitable for individuals who find meditation practice prohibitively difficult.

## Supporting information

Significant features

PC weightings

Correlations between significant features

