## Supplementary Materials for "Uncovering a stability signature of brain dynamics associated with meditation experience using massive time-series feature extraction"

Since our study involves a data-driven approach without hypotheses based on a specific direction of differences within a specific measure from the data, we have not provided hypotheses in our introduction. However, in case it is useful, we have stated our a priori expectations in the following sentences. Based on a conservative view of the degree of difference in neural activity between meditators and non-meditators, we expected that the groups would be classified with modest prediction accuracy, reflecting the large inter-individual variability in both brain activity and the effects of meditation on the brain. Furthermore, since a feature set as comprehensive as that provided by *hctsa* has not been used in any previous research, we also expected that we would obtain a novel list of features which would differentiate the two groups (without specific directional hypotheses).

**Procedure**

Participants in the first study completed the following tasks in sequential order: a Go/No-Go task, a colour Stroop task, an emotional Stroop task, a 2-back working memory task with a dual task tactile distractor, and a Sternberg working memory task. Participants in the second study completed the following tasks in sequential order: a Go/No-Go task, an auditory oddball task, an attention blink task, and had EEG recorded while combined with Transcranial Magnetic Stimulation (TMS). All participants had resting-state EEG data recorded before the TMS was applied, so the application of TMS did not affect the results of our study.

**EEG Pre-processing**

An automatic EEG cleaning toolbox (RELAX) was used to clean the EEG data of artifacts while also preserving neural activity (Bailey et al., 2022a; Bailey et al., 2022b). RELAX implemented the following steps in the following order. Data were high pass filtered within the toolbox using a fourth order Butterworth filter at 1Hz, and a fourth order Butterworth notch filter was applied from 47 to 53Hz. Following PREP's automatic bad electrode detection and removal method (Bigdely-Shamlo et al., 2015), RELAX's default bad electrode detection and removal approach was used. Extreme outlying data periods were also marked for exclusion. Two sequential Multi-channel Wiener Filter (MWF) cleaning steps were applied using a delay period of 16 (so that 32 samples and 32 ms surrounding each timepoint were taken into account when constructing the artifact and clean data templates and when applying the spatial-temporal MWF cleaning to the data). In the first round of

MWF cleaning, muscle activity was reduced. The second round of MWF cleaning addressed any horizontal eye movements that occurred (which can occur even while participants have their eyes closed) and any voltage drift in the data (Somers et al., 2018). To allow for better ICA performance, low pass filtering was then applied to the data at 80 Hz, and PREP's robust average re-referencing was applied (Bigdely-Shamlo, Mullen, Kothe, Su, & Robbins, 2015). cudaICA was used to perform Extended-Infomax Independent Component Analysis on the data, separating the data into its underlying independent components (Raimondo et al., 2012). Components identified as artifacts by ICLabel were then reduced using Wavelet enhanced ICA (Castellanos & Makarov, 2006) before data was reconstructed back into the scalp space (Pion-Tonachini et al., 2019). Removed electrodes were interpolated back into the data using spherical spline interpolation (Perrin et al., 1989).

The data were then split into 30 s epochs (containing 30,000 samples). This epoch length was chosen in the current study as it has successfully been used with the *hctsa* toolbox to categorize sleep stages and predict response to transcranial magnetic stimulation for depression in previous research (Bailey et al., 2023e; Decat et al., 2022). These 30 second epochs were obtained starting from the beginning of the cleaned EEG file, with another 30 s epoch extracted starting from every subsequent 10 s mark throughout the EEG data. Each epoch was then baseline corrected by subtracting the average amplitude across the entire epoch. RELAX's default epoch rejection settings were then applied with the exception that an epoch was only rejected if the voltage shift within the epoch exceeded 120 microvolts. Finally, the first artifact-free 30 s epoch from each participant was selected for inclusion in the analyses. One non-meditator participant's data were excluded at this stage as no clean 30 s epoch could be obtained. This first epoch that remained after each of these steps was downsampled to 160 Hz, which provided a sampling rate sufficiently above the 80 Hz low-pass filter that patterns could be detected even in the highest frequencies remaining in the data after filtering (so that essentially all of the scalp detectable EEG activity would be accounted for by each of the *hctsa* analyses, but none of the features would analyse frequencies that had been filtered out of the data). This meant that the *hctsa* features would be computed on the optimal frequency range. In particular, the *hctsa* measures of oscillations would be focused only on data up to 80 Hz, rather than unnecessarily assessing frequencies above 80 Hz. This is important, as *hctsa* performs the feature computations in a manner that is not informed by the sampling rate of the data, as it is a generic feature set suitable for time-series from all domains (accounting for processes that evolve across a wide range of different timescales). The application of all of these steps resulted in a 64 electrode x 4800 sample multivariate time-series, reflecting a 64 electrode x 30 s extract of the cleaned data.

*hctsa* requires a single time series for the computation of the time-series features. As such, we performed PCA to extract the top components that explained >95% of the variance within the EEG data for inclusion in the *hctsa* computations. This required retaining a total of eight components. Prior to the PCA, each participant's 64 electrode x 4800 sample multivariate time-series data was *z*-transformed (based on all values in the 30 second electrode x timepoint matrix). This preserved potential amplitude relationships between electrodes, while normalising the overall amplitude of each participant's data prior to the PCA (note that each epoch had also been baseline corrected as part of the data pre-processing). The *z*-transform was performed to prevent the potential that imbalances between individuals in EEG amplitude might bias the PCA result towards individuals with larger amplitude EEG data, or towards one of the groups due to an imbalance in variance between the groups. Then, all data was concatenated into a single 64 electrode x 456,000 samples matrix, and a PCA decomposition was performed using MATLAB's **pca** function, which provided a 64 component x 456,000 samples matrix with the first component reflecting a linear combination of the original electrode space time-series that explains the most variance, the second component reflecting a linear combination of the original electrode space time-series that explains the most variance after the variance explained by the first component is removed, and so on for the rest of the components. Concatenating all data prior together to the PCA ensured all participants had the same PCA weights (reflecting spatial maps of weightings from each electrode to each component) applied to their data. All components except for the top eight were then removed, and data from each participant was then separated again, providing an eight-component x 4800 sample time-series data for each individual. PCs 1 through 8 cumulatively explained 95.37% of the variance in the data (see Supplementary Table S1 for the variance explained by each PC). Finally, the first component from all individuals was provided to the *hctsa* feature computation step, then separately the second component from all individuals was provided, and so on until we obtained eight separate individual x feature sets (one individual x feature set for each component).

| Principal Component | Variance Explained (%) |
|---|---|

| | |
|---|---|
| 1 | 50.82 |
| 2 | 19.53 |
| 3 | 13.09 |
| 4 | 4.15 |
| 5 | 3.90 |
| 6 | 1.91 |
| 7 | 1.00 |
| 8 | 0.96 |

Table S1. The amount of variance explained by each of the first eight principal components.

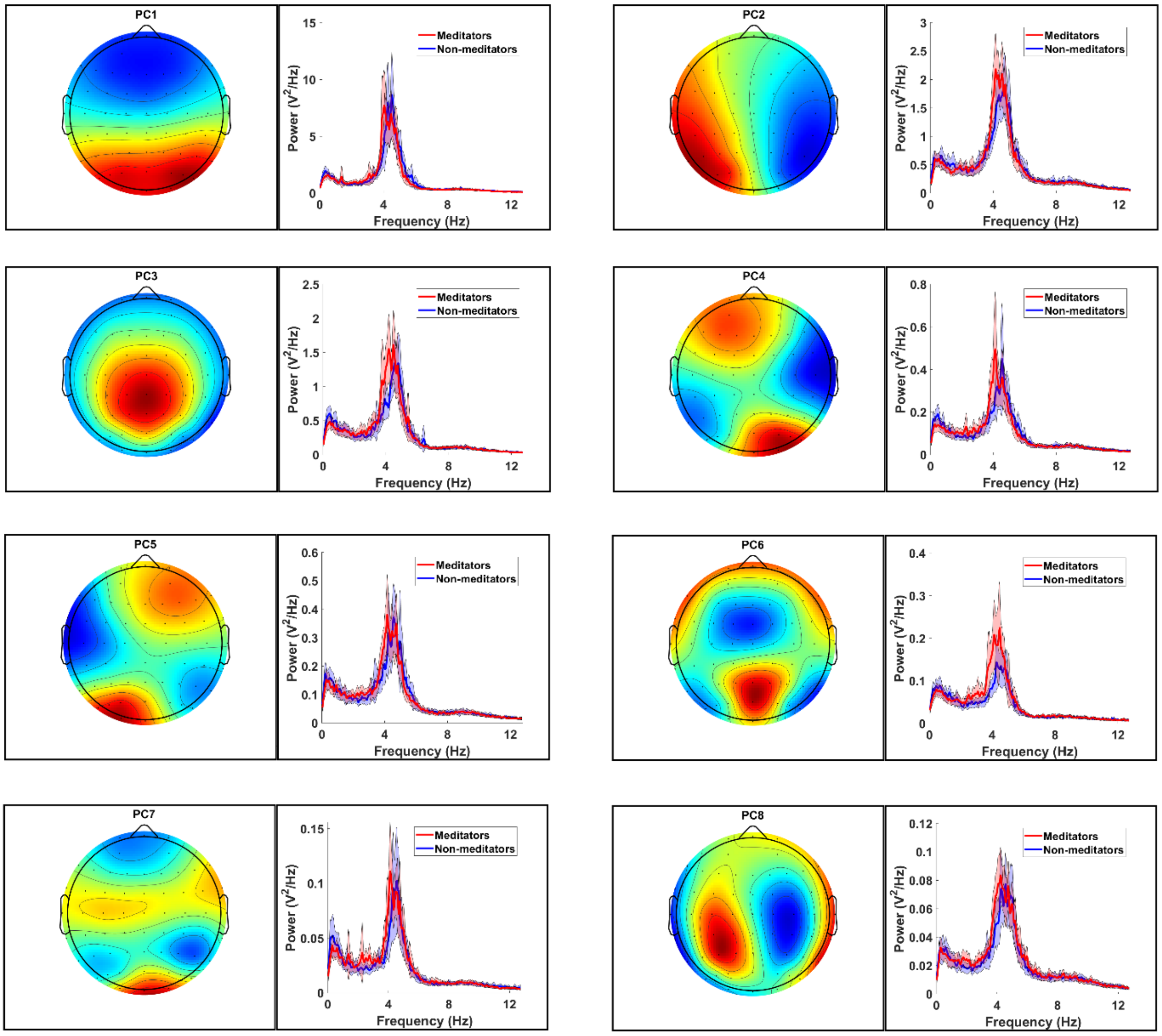

Figure S1. The topography and power spectrum for all principal components. For each PC, on the left is the topographical map of weightings from each electrode that contributed to the PCs. On the right is the mean Welch transformed power spectrum averaged across all participants within each group separately, with the confidence interval shading reflecting 95% confidence intervals.

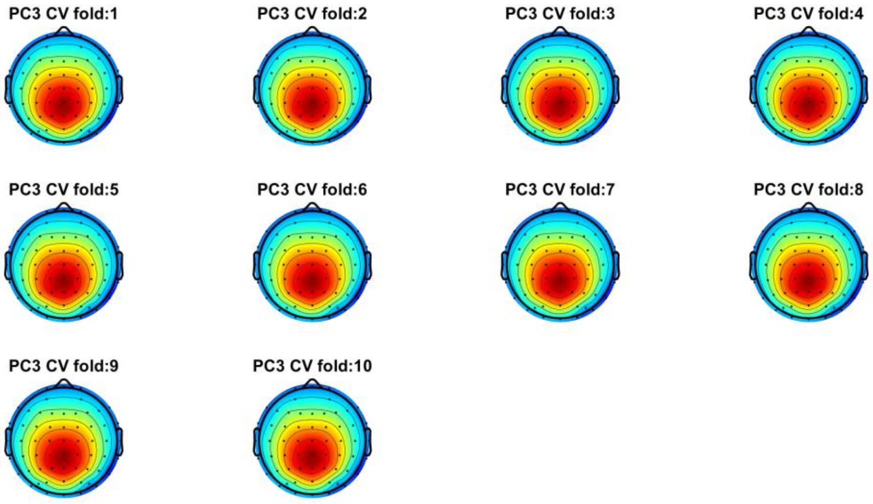


Figure S2. Topographical plots of the weightings provided by PC3 for ten different cross-validation folds (where ~10% participants were excluded at random from the PCA). Note that each PC3 weighting topography is almost identical (the correlation between the weightings of the 10 PCA decompositions were all $r > 0.999$), demonstrating the low probability that including all data in the initial PCA decomposition would have biased the statistical tests.

**Normalisation with the mixed sigmoid transform**

The time-series of all 95 participants remaining in the dataset after the artifact cleaning steps provided more than 80% well-behaved features, so all participants were included in the analysis. Feature normalisation was performed to allow more straightforward comparisons of features measured on different scales and with different distributions by the classifier. We normalised the feature matrix by applying a mixed sigmoid transform to each feature. Mixed sigmoid uses the scaled robust sigmoid to transform the data by default, but reduces to a standard sigmoid when the data has an interquartile range of zero. The scaled robust sigmoid substitutes the typical mean and standard deviation with the median and a multiple of the interquartile range, making it more robust to non-normal distributions (Fulcher et al., 2013). It then linearly rescales the data onto the unit interval.

**Individual Feature Statistics**

To test for differences in individual features, we used the Mann–Whitney U test, as it is a non-parametric test that is robust to violations of normality, making no assumption about any specific distribution (McKnight & Najab, 2010). Due to the computation time required when using the exact Mann–Whitney U test, we first used the approximate test (Fagerland & Sandvik, 2009). Following this, we verified the accuracy of the approximate tests using the exact test for twice the number of features that showed a significant effect using the approximate test, which ensured the exact test accuracy of the number of significant features detected.

The feature named **FC_LocalSimple_median7_sws** in *hctsa*, and referred to as the 'stationarity of median-model predictability feature' for simplicity within this article displayed the strongest ability out of all *hctsa* features at distinguishing meditators from non-meditators ($p_{FDR}$ = 0.012, Cohen's $d$ = 0.911, Figure 4A). As depicted schematically in Figure S2, this feature assessed predictability by computing the median of 7 consecutive samples, then used that median to forecast the next time-series sample. Comparing the standard deviation of the residuals from this prediction process across five non-overlapping segments of the time-series, meditators showed a more consistent level of residual variance across segments than non-meditators (Figure 4A). That is, meditators showed more consistency in the predictability of their dynamics within PC3 than non-meditators.

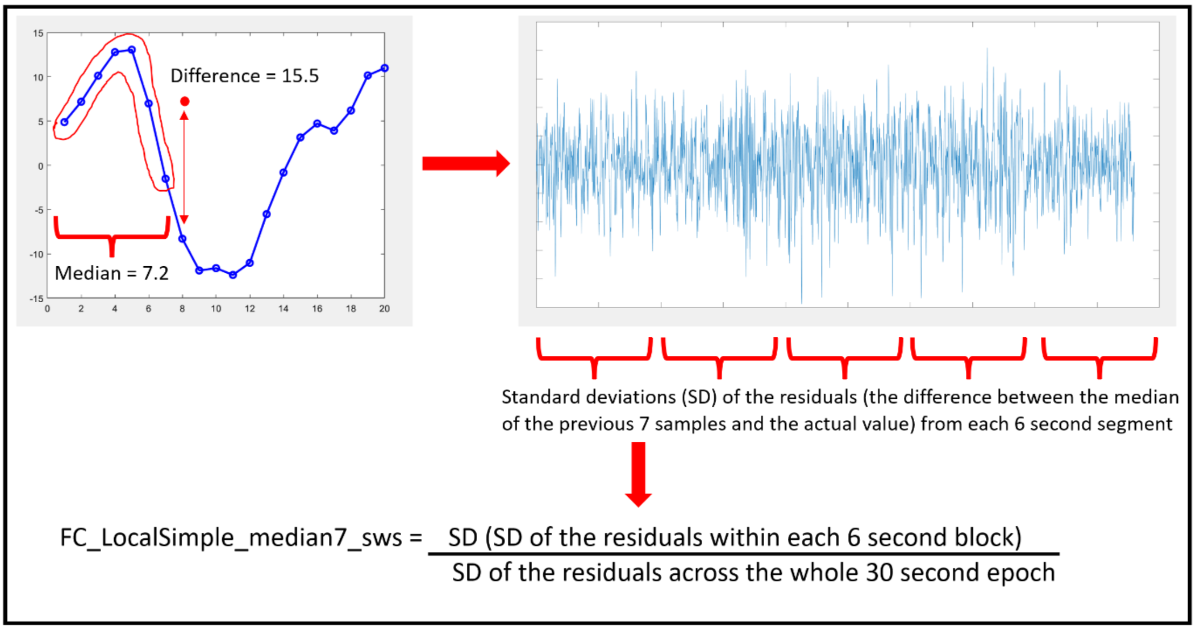


Figure S3. Depiction of the calculation of the top-performing time-series feature, **FC_LocalSimple_median7_sws** ('stationarity of median-model predictability'), which captures the consistency of time-series predictability across different six second segments of the data (which was higher in meditators than non-meditators). This feature showed the largest effect size at differentiating meditators and non-meditators, reflecting the stability of the prediction error within each participant's neural activity from six second segment to six second segment across the 30 second epoch.

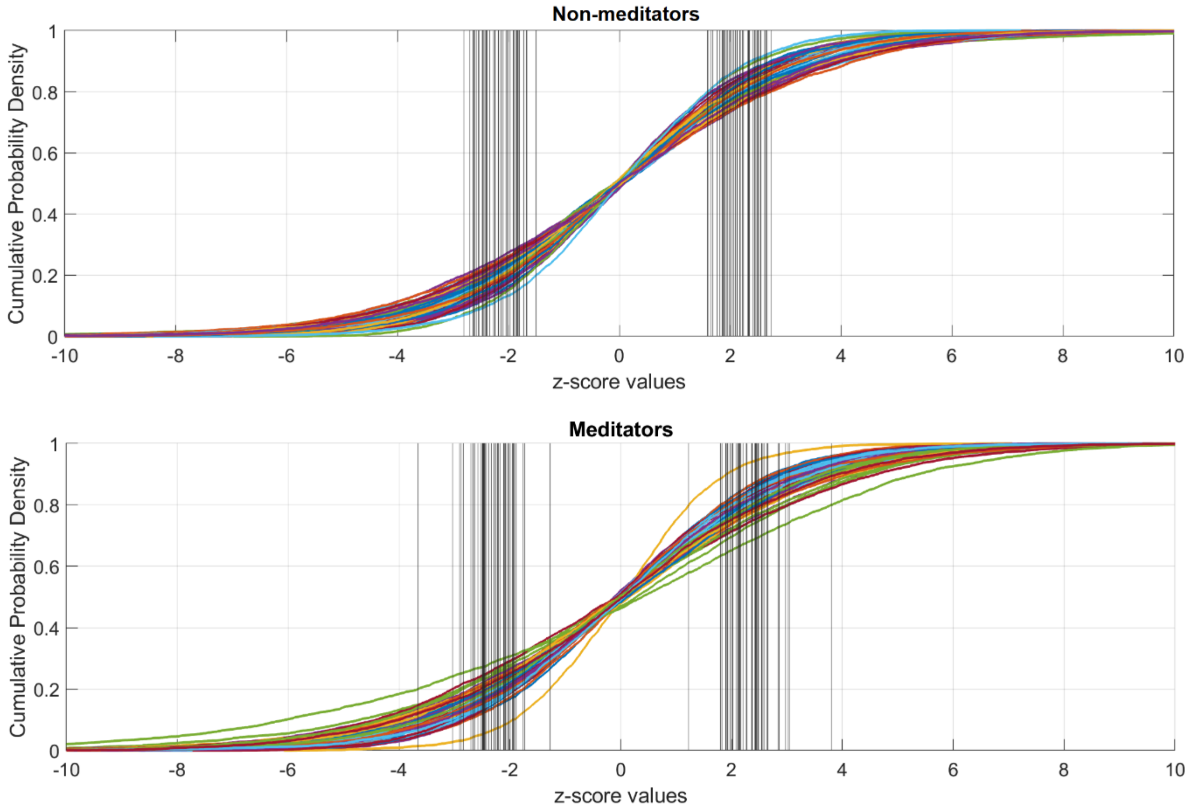


Figure S4. Cumulative density plots showing the distribution of *z*-score values from PC3 for both groups. Black vertical lines are plotted at the 20th and 80th percentiles to visualise the **quantile_20** and **quantile_80** features. On average, the 20th and 80th percentiles are further from the *z*-score 0 point (i.e., the mean) for the meditation group.

**Feature Parameter Exploration**

It is valuable to note that *hctsa* takes an agnostic approach to both the length of the data and the sampling rate and does not endeavour to fine-tune the analysis of features to maximise differentiation between the groups. While *hctsa* tests a remarkably comprehensive set of time-series features, it is not (and could never be) exhaustive. As such, it may be that the computation of different features of the data could be fine-tuned to reveal even greater differences between the groups (and that these measures could more accurately reflect the neural mechanisms of action of mindfulness meditation).

For example, the highest performing feature (stationarity of median-model predictability) assessed the variation across segments of the data in how predictable the data was simply from the median of the preceding 7 samples. *hctsa* assessed the same measure from a range of preceding timepoints from 3, 5, and 7, but did not assess the median from 9 or more preceding timepoints. In order to determine whether variations in the parameters for feature computations might lead to even larger effects, we conducted some additional exploratory tests focused on the 'stationarity of median-model predictability' and 'kurtosis stationarity' features. In an exploratory test of whether different parameter settings for **FC_LocalSimple_medianX_sws** produced different results, we tested medians calculated from 9, 11, 13 and 15 preceding datapoints within PC3, and found that the effect size was even larger when the median was computed from 11 preceding datapoints (Cohen's $d$ = 0.936). While *hctsa* is able to highlight relevant types of time-series properties for differentiating the brain activity of meditators and non-meditators, this result indicates that there is scope for the time-series analyses to be further explored and optimized to provide even greater differentiation of the two groups.

Both the 'stationarity of median-model predictability' and 'kurtosis stationarity' features assessed the stationarity of the data by splitting the time series into segments of equal length (one fifth of the length of the time series, corresponding to 6s in our data) and assessing the variability between those segments. To test the robustness of the 'kurtosis stationarity' feature to different parameter settings, we tested a range of different segment lengths (2 to 15) and the amount to which each segment overlapped with its neighbours (0% to 90%) and found that the effect size was even larger when 14 non-overlapping segments (2.14 seconds per segment) were used (Cohen's $d$ = 1.038). These results indicate that although our approach demonstrated a novel list of features that performed well at differentiating meditators and non-meditators, optimized versions of these features are likely to perform even better. It also suggests that the relevant timescale for the higher kurtosis stationarity shown by the meditators involves data segments in the 2 to 6 second range. It may also be interesting to look at data epochs longer than 30 seconds, to determine whether the same differences in stationarity are revealed when analysed across longer time periods.

Since PC3 contained a prominent alpha peak, we also explored whether the effect detected by the 'stationarity of median-model predictability' feature was driven by the patterns within participant's alpha oscillations. The PC3 time-series data was band-pass filtered so that it just contained alpha activity (using a 4th-order Butterworth filter from 7 to 14 Hz), and the feature was re-computed from the filtered data. The effect size was almost identical (Cohen's $d$ = 0.914). The alpha filtered and original values were also very strongly correlated (Spearman's $\rho = 0.903$, $p < 0.001$). This indicates that the difference between the meditators and non-meditators in the 'stationarity of median-model predictability' feature is likely to be driven by stationarity within participant's alpha oscillations.

Similarly, the effect for the 'kurtosis stationarity' feature was likely to be at least partially driven by kurtosis stationarity within alpha oscillations, as when the data was filtered so that it just contained alpha activity (using a 4th-order bandpass Butterworth filter from 7 to 14 Hz) the difference was still present, but with a smaller effect size (Cohen's $d = 0.704$). The values for the alpha-filtered data also correlated with the original non-filtered values (Spearman's $\rho = 0.699$, $p < 0.001$), but not as strongly as the correlation between the alpha filtered and non-filtered dynamic features, indicating this measure was likely influenced by frequencies other than just alpha oscillations (95% CI for Spearman's $\rho = 0.580$ to 0.790 for the correlation between alpha-filtered and non-filtered 'kurtosis stationarity' compared to 95% CI for Spearman's $\rho = 0.857$ to 0.934 for the correlation between alpha-filtered and non-filtered 'stationarity of median-model predictability').

**Other features that differentiated meditators and non-meditators**

Most features that provided strong differences between the two groups provided a measure of conceptually similar concepts to the three clusters of features reported in our main results, for example distribution shape and stationarity. However, some of these features were also strongly uncorrelated with any other feature, suggesting those features reflect a different aspect of the data from those reported in our results. We have reported a full list of these features in our supplementary materials and provide an illustrative interpretation for one of these features. **ST_MomentCorr_002_02_mean_std_sqrt_density** in *hctsa* ($p_{FDR} = 0.012$, Cohen's $d = 0.808$) first computes a square root transform of the absolute values for all data, then computes the mean and standard deviation in each 600ms segment of the data (with a 20% overlap in each segment), then assesses the range of the mean across all segments multiplied by the range of the standard deviation of values across all segments, divided by the total number of samples. As such, this feature is influenced by both the distributional and stationarity properties of the data, similar to the clusters reported in our main results. For this feature, meditators showed lower values in their PC3 time-series, suggesting smaller extreme variations in their time series data (Figure S5). Combined with the **quantile_20** and **quantile_80** features, these results suggest that the meditator group showed larger variation in their typical EEG amplitude from the weightings captured by PC3, while the non-meditator group showed larger extreme deviations within specific brief

(600ms) time periods. To enable researchers to explore the additional features for potential use in future research, the full list of features is presented in our supplementary materials.

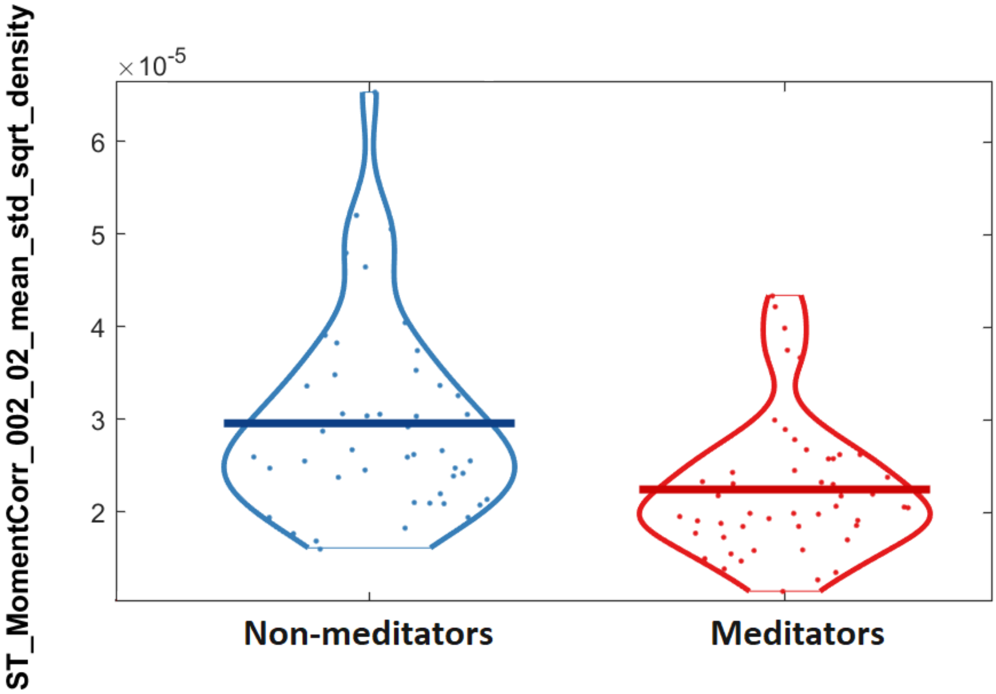


Figure S5. The distribution of values from the **ST_MomentCorr_002_02_mean_std_sqrt_density** feature. This feature assessed the range of the mean multiplied by the standard deviation of the absolute transformed values in each 600ms period of the data, providing high values if any 600ms period across the 30 second epoch contained an extremely high or low mean, or if any of these 600ms period contained large variation. For this feature, meditators showed lower values, suggesting less extreme variations in their time series data ($p_{FDR}$ = 0.012, Cohen's d = 0.808).

We note that other features also performed highly, including the smaller clusters of features that appeared within the set of the top 50 features (Figure S6). Additionally, features that showed the top 51 to 405 strongest effects also provided statistically significant differentiation of meditators and non-meditators. Most of these features provided measures of similar concepts to the top 50 features, both correlated strongly and clustering with these top 50 features. However, some of these additional features were not related to our top three clusters, which capture: (i) stationarity in dynamic properties; (ii) stationarity in distributional properties; and

(iii) global measures of distributional shape. As such, these lower-performing (but still statistically significant) features may provide further novel insight into the difference in brain activity between meditators and non-meditators. Our supplementary materials contain a full list of significant features from PC3 and a corresponding feature–feature correlation matrix.

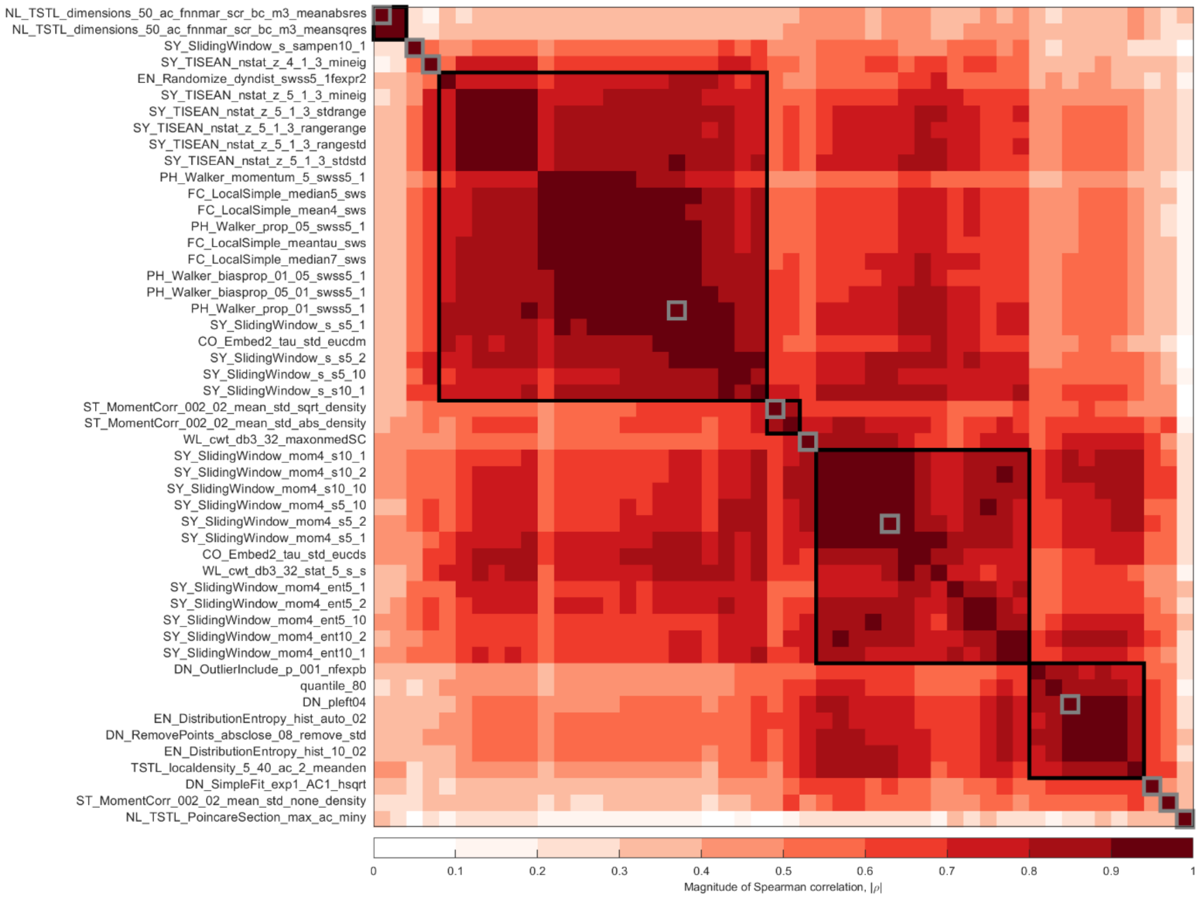


Figure S6. The cluster plot of the top 50 features from PC3 labelled using *hctsa* labels (which are available for interpretation within the toolbox). Features are clustered using absolute Spearman correlations and a threshold of |ρ| = 0.75 for forming clusters, enabling visualisation of how strongly the different clusters are related to each other.

**Confirmatory Analysis Using PCA Decompositions with Unseen Test Set Data**

To confirm that the inclusion of our test set data in our PCA decomposition did not bias our results, we performed 10 separate PCA decompositions, each of which excluded approximately 10% of the (randomly selected) participants. We then applied the weightings for PC3 from each of these training PCA decompositions to only the excluded (test set) participants, such that the PC3 time-series obtained for each participant was produced by a PCA decomposition that did not include the participant. We then re-computed the individual feature values from the PC3 time-series for each participant for the three highest performing representative features of each cluster that our primary analysis showed to differentiate the two groups (**SY_SlidingWindow_mom4_ent5_10, FC_LocalSimple_median7_sws**, and **quantile_80**). We then used the Mann-Whitney U test to analyse the differences between groups for these three features. Once we obtained the uncontrolled *p*-value from this process, we replaced the matching individual feature p-values from our primary analysis of these features with these new *p*-values. To control for multiple comparisons, we then applied the same FDR correction to the 7,381 *p*-values. The results of these analyses showed that these features still significantly differentiated the meditator and non-meditator groups, with the result in the same direction as our primary analysis (**SY_SlidingWindow_mom4_ent5_10:** $p_{FDR}$ = 0.0169, Cohen's *d* = 0.938**,** **FC_LocalSimple_median7_sws:** $p_{FDR}$ = 0.0169, Cohen's *d* = 0.902, and **quantile_80:** $p_{FDR}$ = 0.0205, Cohen's *d* = 0.814). Finally, we used a linear support-vector machine (SVM) to classify meditators and non-meditators based on these features and used 10-fold cross-validation to compute the balanced accuracy from this analysis, which was compared with a model-based null permutation with 1000 null samples. The results showed a mean balanced accuracy of 74.01%, which significantly exceeded chance against the model-based null permutation test (p = 0.005). This replicates the results from our primary analyses reported in the manuscript. However, it is worth noting that while this exploratory analysis was performed to verify our primary comparisons, and that the result performed better than a model-based null trained on the same features, conducting the statistical test on individual features selected based on their ability to differentiate the groups in the same dataset is likely to bias the results (as the features were selected dependent on their ability to differentiate the two groups). As such, this test of classification accuracy is not valid as more than a confirmation of our primary analysis. The accuracy of a classification model that includes only these three features should be tested on an independent dataset to provide a valid test of the classification accuracy of these features.

**Supplementary Discussion Points**

***The advantages of a comprehensive analysis data-driven approach over hypothesis driven approaches***

Our findings that features assessing data stationarity within PC3 provided the largest effect sizes, combined with the considerable focus of previous research on oscillations and the almost complete absence of measures assessing stationarity in previous research highlights the valuable and novel contribution of using *hctsa* (or another comprehensive data driven approach) to determine how neural activity differs in meditation. Our perspective is that this approach revealed features of the data that differentiated meditators and controls which would never have been tested using a hypothesis driven approach. By way of example, the feature that strongest effect size was **FC_LocalSimple_median7_sws**. This feature measures the variability in time-series predictability across five (6 s-long) non-overlapping segments of the 30 s PC3 time series. As depicted schematically in Figure 5, this feature assesses predictability by computing the median of 7 consecutive samples and uses that median to forecast the next time-series sample. It then computed the standard deviation of the residuals from this prediction process across five non-overlapping segments of the time-series. Figure 4 depicts this calculation visually. Essentially, this feature answers the question "across the 6s blocks, does the variability in prediction misses vary more than the variability in prediction misses across the whole 30s?", providing a measure of how stationary the variability in predictability is across the time series. Meditators showed lower values, indicating that the variability in how accurately their data could be predicted did not change from 6 second block to 6 second block across the epoch. We hope this explanation provides the reader with a good understanding of how unlikely it would be for a researcher to generate a hypothesis that the two groups would differ in this feature of the data via theoretical understanding.

It is also worth noting that the features from the stationarity and distributional properties clusters were correlated with features from other clusters, suggesting the different clusters did not assess entirely independent characteristics of the data. As such, the features that were best at distinguishing the two groups were likely to be influenced by non-independent mechanisms within our dataset. However, the separate clusters of features also captured some independent variability from each other, with the features measuring the distribution of PC3 and features measuring the stationarity of the dynamical properties of PC3 capturing more independent variability from each other.

***A potential neurofeedback application based on our results***

With regards to the potential application of neurofeedback based on the stationarity features - in the case that the kurtosis stationarity feature from our study is validated by future research, a target amount of kurtosis from PC3 within short (~1 second) time periods could be established, and individuals could be provided with positive feedback as neural activity detected within PC3 converges with this target amount. Long-term repetition of this practice might improve the ability of these individuals to generate this consistent level of kurtosis, potentially leading to more stable neural activity within PC3, and brain activity more similar to the pattern shown by the experienced meditators in our study. If this neurofeedback approach is successful, it might be associated with similar well-being and attention benefits to those reported by and detected in experienced meditators. However, it is worth noting that valid estimation of kurtosis requires a high number of datapoints (Bai & Ng, 2005), so high sampling rates may be required, and future work may need to test the timescale that balances accurate estimation and sufficiently rapid feedback (alternatively, the 'stationarity of median-model predictability' measure "**FC_LocalSimple_median7_sws**" may be more practical to apply as a potential neurofeedback target). It is also possible that "device-oriented" approaches like this may be a distraction from a meditation practice that may already be optimal when taught within certain traditions, so it may be that this proposed neurofeedback approach would be more suitable for individuals who find meditation practice prohibitively difficult.

***Additional Limitations***

One additional limitation to our conclusions is that although participants were instructed to "rest, not meditate", it is not possible to confirm that participants were resting rather than meditating. It may even be that many meditators naturally rest in a somewhat meditative state. As such, it is not possible to confirm that our results reflect-state differences in brain activity in meditators rather than meditation-state related brain activity. Finally, we note that while many individual features provided strong effect sizes for the difference between the two groups, there was still considerable overlap in the values from the meditator and non-meditator groups within all features, such that no individual feature could identify an individual as a meditator with 100% accuracy. This is not unexpected given the likely diversity of neural activity patterns across any two populations, across the range of meditation types and experience, and within typically noisy EEG data.

**Supplementary Materials References**


Bai, J., & Ng, S. (2005). Tests for skewness, kurtosis, and normality for time series data. *Journal of Business & Economic Statistics*, *23*(1), 49-60.

Bailey, N., Biabani, M., Hill, A. T., Miljevic, A., Rogasch, N. C., McQueen, B., Murphy, O. W., & Fitzgerald, P. (2022a). Introducing RELAX (the Reduction of Electroencephalographic Artifacts): A fully automated pre-processing pipeline for cleaning EEG data-Part 1: Algorithm and Application to Oscillations. *bioRxiv*. https://doi.org/https://doi.org/10.1101/2022.03.08.483548

Bailey, N., Fulcher, B., Arns, M., Fitzgerald, P. B., Fitzgibbon, B., & van Dijk, H. (2023e). Prediction of response to transcranial magnetic stimulation treatment for depression using electroencephalography and statistical learning methods, including an out-of-sample validation. *Open Science Framework*. https://doi.org/osf.io/pexgn

Bailey, N., Hill, A., Biabani, M., Murphy, O., Rogasch, N., McQueen, B., Miljevic, A., & Fitzgerald, P. (2022b). Introducing RELAX (the Reduction of Electroencephalographic Artifacts): A fully automated pre-processing pipeline for cleaning EEG data – Part 2: Application to Event-Related Potentials. *bioRxiv*. https://doi.org/https://doi.org/10.1101/2022.03.08.483554

Bigdely-Shamlo, N., Mullen, T., Kothe, C., Su, K.-M., & Robbins, K. A. (2015). The PREP pipeline: standardized preprocessing for large-scale EEG analysis. *Frontiers in Neuroinformatics*, *9*, 16.

Castellanos, N. P., & Makarov, V. A. (2006). Recovering EEG brain signals: Artifact suppression with wavelet enhanced independent component analysis. *Journal of Neuroscience Methods*, *158*(2), 300-312.

Decat, N., Walter, J., Koh, Z. H., Sribanditmongkol, P., Fulcher, B. D., Windt, J. M., Andrillon, T., & Tsuchiya, N. (2022). Beyond traditional visual sleep scoring: massive feature extraction and unsupervised clustering of sleep time series. *Sleep Medicine*, *98*, 39-52.

Fagerland, M. W., & Sandvik, L. (2009). The wilcoxon–mann–whitney test under scrutiny. *Statistics in medicine*, *28*(10), 1487-1497.

Fulcher, B. D., Little, M. A., & Jones, N. S. (2013). Highly comparative time-series analysis: the empirical structure of time series and their methods. *Journal of the Royal Society Interface*, *10*(83), 20130048.

McKnight, P. E., & Najab, J. (2010). Mann-Whitney U Test. *The Corsini encyclopedia of psychology*, 1-1.

Perrin, F., Pernier, J., Bertrand, O., & Echallier, J. F. (1989). Spherical splines for scalp potential and current density mapping. *Electroencephalography and clinical neurophysiology*, *72*(2), 184-187.

Pion-Tonachini, L., Kreutz-Delgado, K., & Makeig, S. (2019). The ICLabel dataset of electroencephalographic (EEG) independent component (IC) features. *Data in brief*, *25*, 104101.

Raimondo, F., Kamienkowski, J. E., Sigman, M., & Fernandez Slezak, D. (2012). CUDAICA: GPU optimization of infomax-ICA EEG analysis. *Computational intelligence and neuroscience*, *2012*.

Somers, B., Francart, T., & Bertrand, A. (2018). A generic EEG artifact removal algorithm based on the multi-channel Wiener filter. *Journal of neural engineering*, *15*(3), 036007. https://doi.org/10.1088/1741-2552/aaac92
